# Overexpression of the RNA-binding protein NrdA affects global gene expression and secondary metabolism in *Aspergillus* species

**DOI:** 10.1101/2021.03.15.435561

**Authors:** Chihiro Kadooka, Kosuke Izumitsu, Teigo Asai, Kazuki Mori, Kayu Okutsu, Yumiko Yoshizaki, Kazunori Takamine, Masatoshi Goto, Hisanori Tamaki, Taiki Futagami

## Abstract

RNA-binding protein Nrd1 plays a role in RNA polymerase II transcription termination. In this study, we showed that the orthologous NrdA is important in global mRNA expression and secondary metabolism in *Aspergillus* species. We constructed an *nrdA* conditional expression strain using the Tet-On system in *Aspergillus luchuenesis* mut. *kawachii*. Downregulation of *nrdA* caused a severe growth defect, indicating that NrdA is essential for the proliferation of *A. kawachii*. Parallel RNA-sequencing and RNA immunoprecipitation-sequencing analysis identified potential NrdA-interacting transcripts, corresponding to 32% of the predicted protein coding genes of *A. kawachii*. Subsequent gene ontology analysis suggested that overexpression of NrdA affects the production of secondary metabolites. To clarify this, we constructed NrdA-overexpressing strains of *Aspergillus nidulans*, *Aspergillus fumigatus*, and *Aspergillus oryzae*. Overexpression of NrdA reduced the production of sterigmatocystin and penicillin in *A. nidulans*, as well as that of helvolic acid and pyripyropene A in *A. fumigatus*. Moreover, it increased the production of kojic acid and reduced production of penicillin in *A. oryzae*. These effects were accompanied by almost consistent transcriptional changes in the relevant genes. Collectively, these results suggest that NrdA is the essential RNA-binding protein, which plays a significant role in global gene expression and secondary metabolism in *Aspergillus* species.

**IMPORTANCE:** Nrd1, a component of the Nrd1–Nab3–Sen1 complex, is an essential RNA-binding protein involved in transcriptional termination in yeast. However, its role in filamentous fungi has not been studied. In this study, we characterized an orthologous NrdA in the *Aspergillus* species, identified potential NrdA-interacting mRNA, and investigated the effect of overexpression of NrdA on mRNA expression in *Aspergillus luchuensis* mut. *kawachii*. The results indicated that NrdA controls global gene expression involved in versatile metabolic pathways, including the secondary metabolic process. We demonstrated that NrdA overexpression significantly affected the production of secondary metabolites in *Aspergillus nidulans*, *Aspergillus oryzae*, and *Aspergillus fumigatus*. Our findings are of importance to the fungal research community because the secondary metabolism is an industrially and clinically important aspect for the *Aspergillus* species.

## INTRODUCTION

There are at least two pathways for transcription termination in the budding yeast *Saccharomyces cerevisiae*, namely the RNA polymerase II cleavage/polyadenylation factor-related pathway and the Nrd1–Nab3–Sen1 (NNS)-related pathway (1, 2). The former cleavage/polyadenylation factor pathway is required for polyadenylation of the 3’ end of precursor mRNA. The latter pathway is required for the early termination of transcripts to produce the shorter polyadenylated transcripts. Moreover, it is involved in the subsequent production of non-coding transcripts, such as small nucleolar RNAs, small nuclear RNAs, cryptic unstable transcripts, and stable uncharacterized transcripts with largely unknown functions. The Nrd1 and Nab3 are essential RNA-binding proteins and bind specific sequences in target RNA, whereas Sen1 is a helicase responsible for the termination of transcription. The NNS complex has been extensively studied from the view point of non-coding RNA production. However, it has been recently suggested that the NNS-related pathway also plays a significant role in protein-coding mRNA involved in response to nutrient starvation (3, 4). It was proposed that binding of Nrd1 and Nab3 led to premature transcription termination, resulting in the downregulation of mRNA levels. In *S. cerevisiae*, it was estimated that Nrd1 and Nab3 directly regulate a variety of protein-coding genes, representing 20%–30% of protein-coding transcripts (5). In addition, the Nrd1 ortholog Seb1 drives transcription termination for both protein-coding and non-coding sequences in the fission yeast *Schizosaccharomyces pombe* (6, 7).

We have investigated the mechanism of citric acid accumulation by the white koji fungus *Aspergillus luchuensis* mut. *kawachii*, which is used for the production of shochu, a Japanese traditional distilled spirit. *A. kawachii* is used as a producer of starch-degrading enzymes, such as α-amylase and glucoamylase (8, 9). In addition, *A. kawachii* also produces a large amount of citric acid, which prevents the growth of microbial contaminants during the fermentation process (10). During the course of the study, we found that the Nrd1 ortholog encoding *nrdA* was located in a syntenic region together with mitochondrial citrate synthase-encoding *citA* and mitochondrial citrate transporter encoding *yhmA* in the subdivision Pezizomycotina (including genus *Aspergillus*). However, it was not found in the subdivisions Saccharomycotina (including genus *Saccharomyces*) and Taphrinomycotina (including genus *Schizosaccharomyces*) (11). Considering the numerous examples of metabolic gene clusters in fungi and plants (12, 13), this finding motivated us to study the function of NrdA in *Aspergillus* species in terms of metabolic activity.

In this study, we showed that NrdA is an essential RNA-binding protein involved in the production of citric acid in *A. kawachii* using the *nrdA* conditional expression strain. Nevertheless, whether *nrdA* is functionally related to *citA* and *yhmA* remains unclear. By combining RNA-sequencing (RNA-seq) and RNA immunoprecipitation-sequencing (RIP-seq) analyses, we identified potential NrdA– mRNA interactions, corresponding to approximately 32% of protein-coding genes of the *A. kawachii* genome. In addition, we showed that overexpression of *nrdA* causes significant changes in the expression levels of NrdA-interacting mRNA involved in secondary metabolism. Furthermore, we demonstrated that overexpression of *nrdA* also caused changes in the production of secondary metabolites in *Aspergillus nidulans*, *Aspergillus fumigatus*, and *Aspergillus oryzae*. These results suggested that the NrdA-associated transcription termination pathway potentially regulates the secondary metabolic process in the *Aspergillus* species.

## RESULTS AND DISCUSSION

### Sequence feature of NrdA

The *A. kawachii nrdA* gene encodes a protein composed of 742 amino acid residues. BLASTP analysis (https://blast.ncbi.nlm.nih.gov/Blast.cgi) showed that the amino acid sequence percent identities between *A. kawachii* NrdA and *S. cerevisiae* Nrd1, and between *A. kawachii* NrdA and *Schiz. pombe* Seb1 were 48.57% and 34.52%, respectively. The RNA polymerase II C-terminal domain-interacting domain (CID) (1–150 amino acid residues of *S. cerevisiae* Nrd1) (14–16) and RNA recognition motif (RRM) (339–407 amino acid residues of *S. cerevisiae* Nrd1) (16, 17) are well conserved in *A. kawachii* NrdA (Fig. S1). The Pfam domain analysis (https://pfam.xfam.org/) also confirmed the presence of CID and RRM in the relevant regions of *A. kawachii* NrdA (data not shown). On the other hand, the amino acid residues of the Nab3-binding domain, arginine-glutamate/arginine-serine-rich domain, and the C-terminal proline/glutamine (P/Q)-rich low-complexity domain (LCD) (151–214, 245–265, and 513–575 amino acid residues of *S. cerevisiae* Nrd1, respectively) (16, 18, 19) were less conserved in the *A. kawachii* NrdA. The Nab3-binding domain, arginine-glutamate/arginine-serine-rich domain, and P/Q rich LCD of *A. kawachii* NrdA were more similar to those of *Schiz. pombe* Seb1 than those of *S. cerevisiae* Nrd1. In addition, amino acid restudies between RRM and P/Q rich LCD were well conserved among *S. cerevisiae* Nrd1, *Schiz. pombe* Seb1, and *A. kawachii* NrdA. Phylogenetic analysis supported that Nrd1 orthologs of subdivision Pezizomycotina (including *A. kawachii* NrdA) are closer to those of subdivision Taphrinomycotina (including *Schiz. pombe* Seb1) than those of subdivision Saccharomycotina (including *S. cerevisiae* Nrd1) (Fig. S2).

### Phenotype of the Tet-*nrdA* strain

To investigate the physiologic role of *nrdA*, we attempted to construct *A. kawachii nrdA* disruptant. However, all of transformants obtained by the *nrdA* disruption cassette were heterokaryotic gene disruptants (data not shown). Therefore, we constructed a Tet-*nrdA* strain that conditionally expressed *nrdA* under the control of a Tet-On promoter. The Tet-*nrdA* strain exhibited significantly deficient growth in yeast extract sucrose (YES) agar medium without doxycycline (Dox) (*nrdA* depletion condition) (Fig. 1A), indicating that disruption of *nrdA* induces lethality. This result was consistent with those of previous reports stating that *NRD1* and *seb1* are essential genes in *S. cerevisiae* and *Schiz. pombe*, respectively (20, 21).

**FIG 1.**
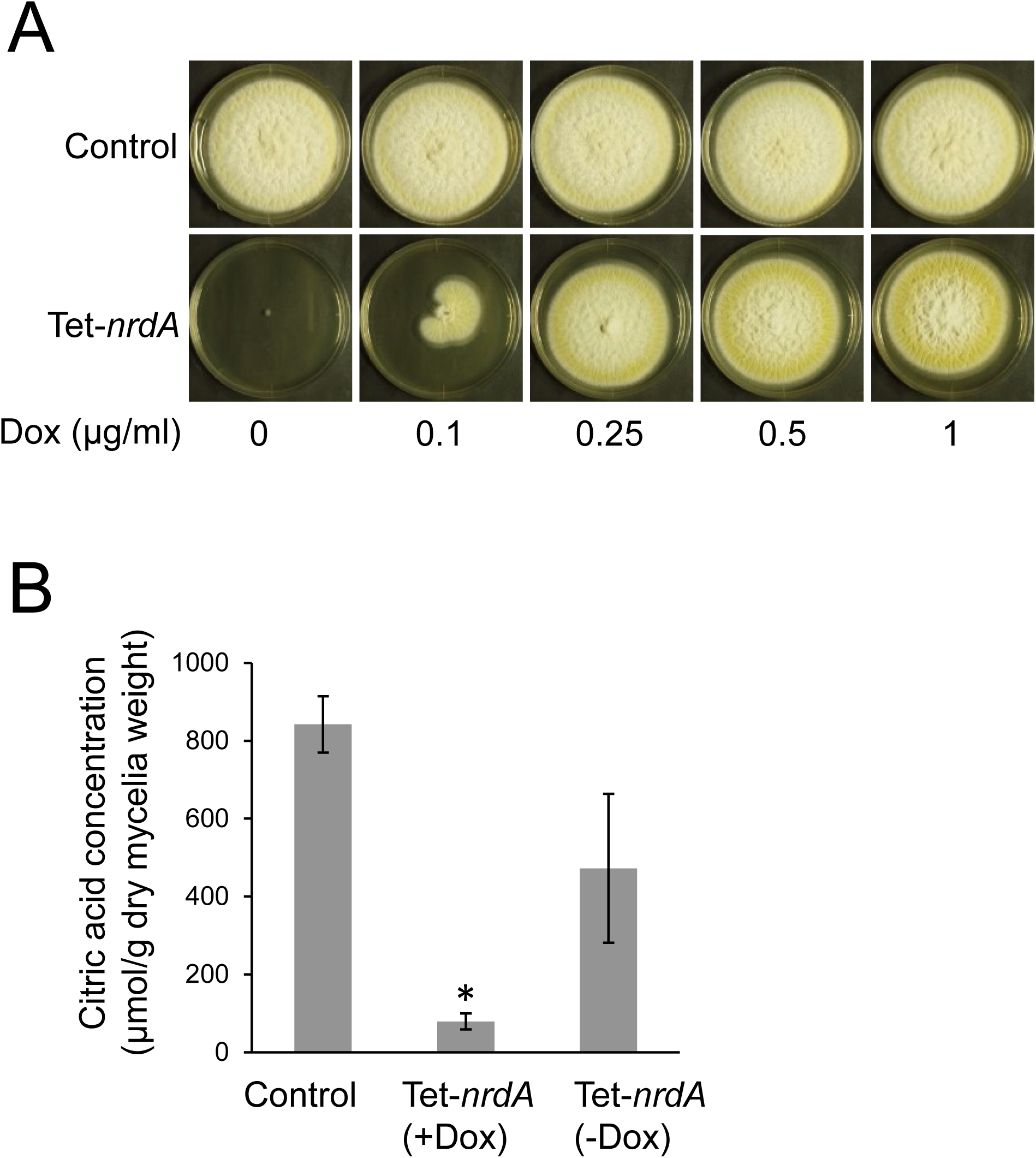
(A) Colony formation of *A. kawachii* control and Tet-*nrdA* strains. Conidia (10^4^) were inoculated onto YES medium. Strains were cultured at 30°C for 5 days on YES medium with or without doxycycline (Dox). Conidia of each strain (1 × 10^4^) were inoculated onto agar medium. (B) Citric acid production of control and Tet-*nrdA* strains. Strains were precultured in YES medium with 1 µg/ml Dox for 16 h, transferred to CAP medium, and further cultivated for 48 h. The mean and standard deviation were determined from the results of 3 independent cultivations. *, statistically significant difference (*p* < 0.05, Welch’s *t*-test) relative to the data obtained for the control strain.

Next, we tested the effect of NrdA expression on citric acid production, because *nrdA* was located in the syntenic region with mitochondrial citrate synthase encoding *citA* and citrate transporter encoding *yhmA* (11). We compared the production of citric acid by *A. kawachii* control and Tet-*nrdA* strains (Fig. 1B). The control strain was precultivated in YES medium at 30°C for 16 h, transferred to citric acid production (CAP) medium (an optimized medium for CAP) (11), and further cultured at 30°C for 48 h. Because the Tet-*nrdA* strain showed severe defect growth in the absence of Dox, it was precultivated in YES medium with Dox and transferred to CAP medium with or without Dox. Following cultivation, the concentration of citric acid in the culture supernatant and mycelial biomass was measured to determine the extracellular CAP per mycelial weight. Based on the level of citric acid in the culture supernatant and amount of mycelial biomass produced, the Tet-*nrdA* strain cultivated without Dox showed similar CAP compared with the control strain. In contrast, the Tet-*nrdA* strain cultivated with Dox exhibited only approximately 9% of the CAP of the control strain.

### Subcellular localization of NrdA

To analyze the subcellular localization of NrdA in *A. kawachii*, NrdA tagged with green fluorescent protein (GFP-NrdA) and histone H2B tagged with monomeric red fluorescent protein (mRFP) (H2B-mRFP, a nuclear marker protein) were co-expressed in the Tet-*nrdA* strain. The GFP-NrdA was expressed by the native *nrdA* promoter. Functional expression of GFP-NrdA was confirmed by the viable phenotype of the Tet-*nrdA* plus *GFP-nrdA* plus *H2B-mRFP* strain in YES agar medium without Dox (Fig. 2A). The Tet-*nrdA* plus *GFP-nrdA* plus *H2B-mRFP* strain was cultivated in minimal liquid medium for 16 h without shaking and observed by fluorescence microscopy. Green fluorescence associated with GFP-NrdA merged with the red fluorescence of H2B-mRFP (Fig. 2B), indicating that the GFP-NrdA localizes in the nucleus. In addition, speckles of green fluorescence were observed in the nucleus. It was reported that the Nrd1-dependent nuclear speckles appear in *S. cerevisiae* under glucose starvation conditions (3). Thus, it might be possible that NrdA also forms these speckles in *A. kawachii* in response to some environmental stress.

**FIG 2.**
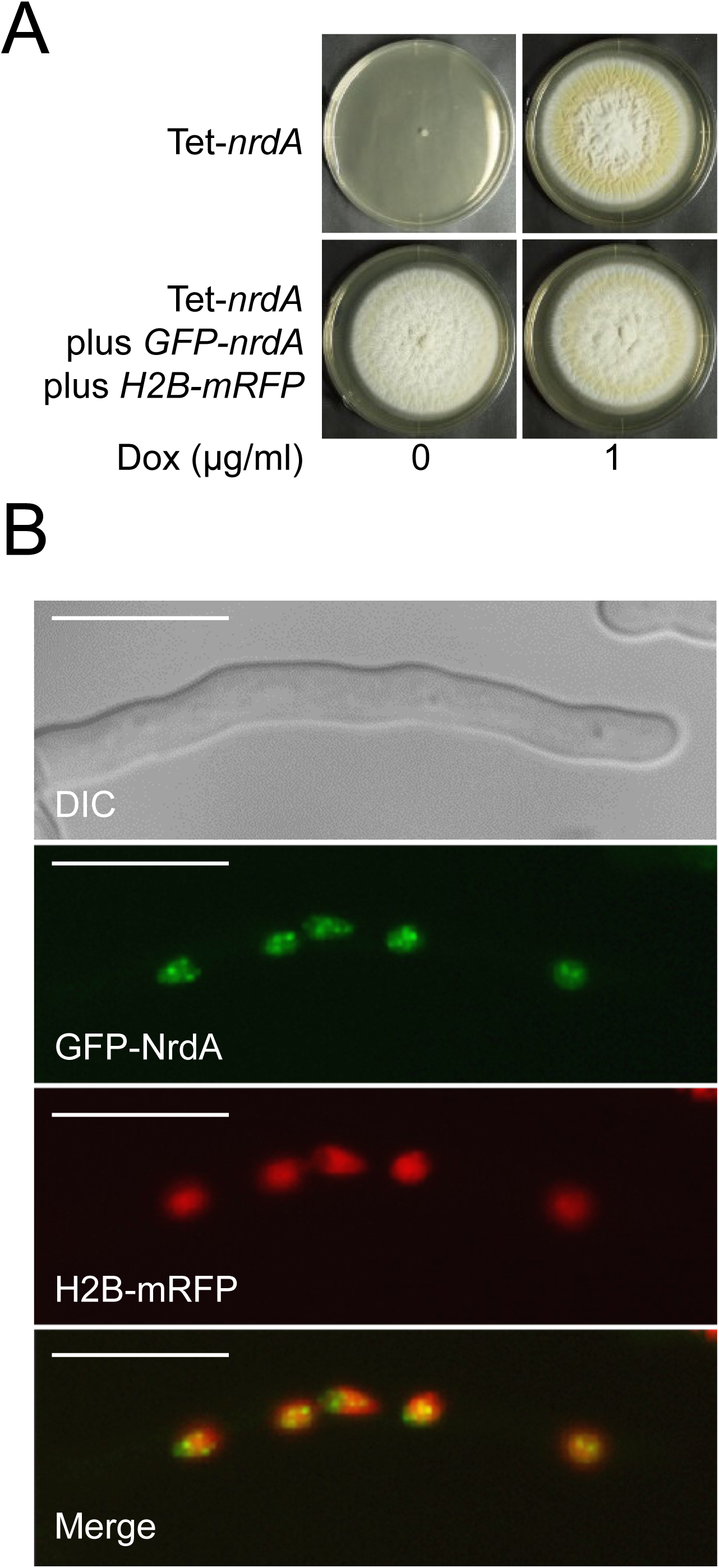
(A) Colony formation of *A. kawachii* Tet-*nrdA* strain and its complemented strain with GFP-NrdA and H2B-mRFP. Conidia (1 × 10^4^) were inoculated onto YES agar medium with or without 1µg/ml Dox and incubated at 30°C for 5 days. (B) Fluorescence microscopic observation of the GFP-NrdA- and H2B-mRFP-expressing strain. Scale bars indicate 10 μm.

### Examination of experimental conditions for transcriptome analysis

We performed transcriptome analysis to investigate the NrdA-associated transcripts. For this purpose, we constructed a Tet-*S*-*nrdA* strain, which expresses S-tagged NrdA (S-NrdA) under the control of the Tet-On promoter. The Tet-*S*-*nrdA* strain formed colonies in the YES agar medium with Dox, whereas it showed severe growth defect in the YES agar medium without Dox. These findings confirmed that the Tet-On system regulates the functional S-NrdA (Fig. 3A). In addition, the expression of S-NrdA was confirmed by the detection of a band of the predicted size of the S-NrdA protein through S-protein affinity purification (Fig. 3B left panel) and detection by immunoblotting using an anti-S-tag antibody (Fig. 3B right panel).

**FIG 3.**
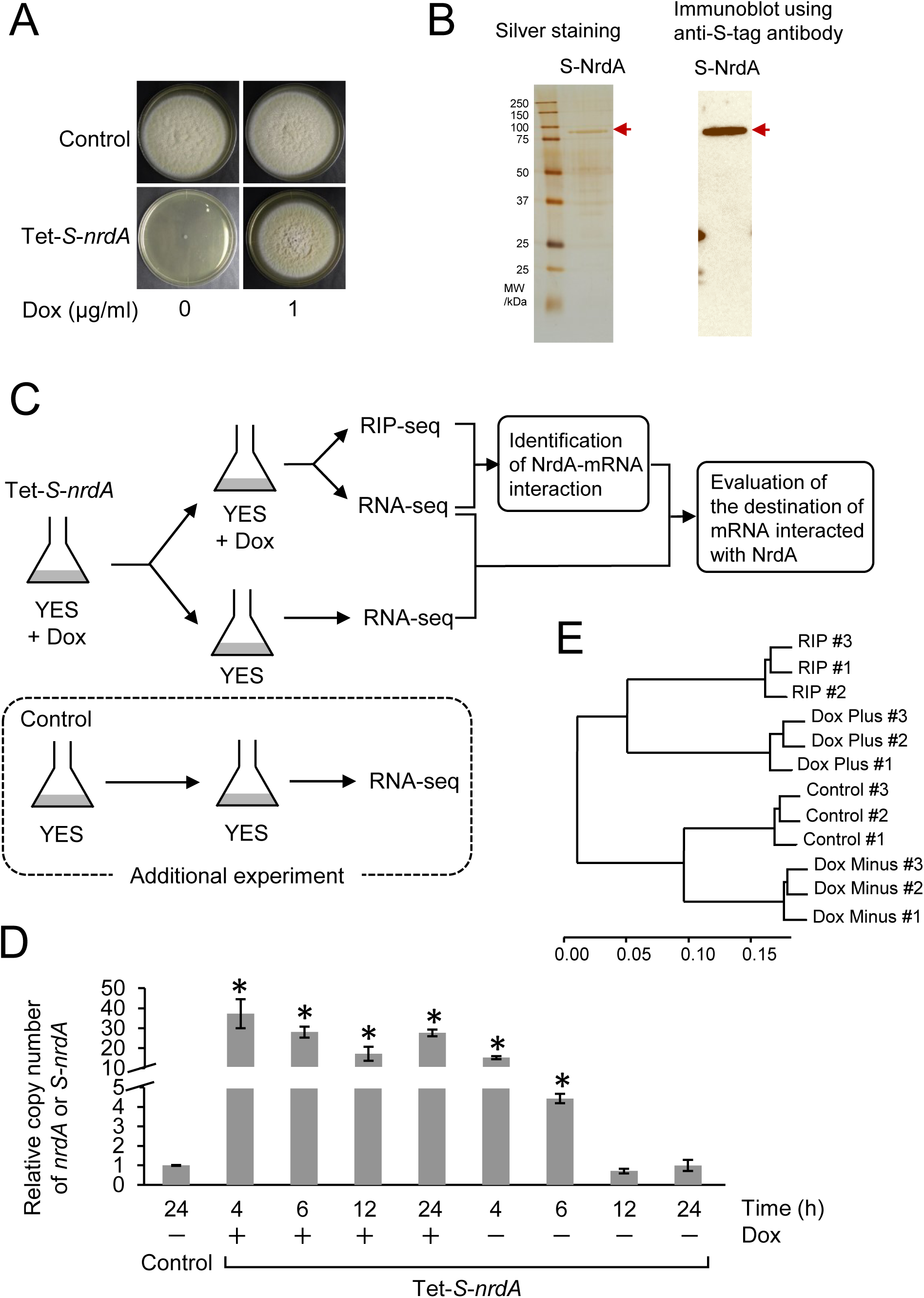
(A) Colony formation of *A. kawachii* control and Tet-*S*-*nrdA* strains. Strains were cultured at 30°C for 5 days on YES medium with or without Dox. Conidia of each strain (1 × 10^4^) were inoculated onto agar medium. (B) Immunoblotting analysis of purified S-NrdA protein from the *A. kawachii* Tet-*S*-*nrdA* strain. The apparent molecular mass of S-NrdA was 80.7 kDa. (C) Scheme of experimental design for the RNA-seq and RIP-seq analyses to identify the NrdA-interacting transcripts and investigate the effect of NrdA overexpression. (D) Quantitative RT-PCR analysis to evaluate the expression levels of *nrdA* and *S*-*nrdA* in *A. kawachii* control and Tet-*S*-*nrdA* strains. (E) Cluster analysis to confirm the *S*-*nrdA* expression condition in the *A. kawachii* Tet-*S*-*nrdA* strain in the presence or absence of Dox. The mean and standard deviation were determined from the results of 3 independent cultivations. *, statistically significant difference (*p* < 0.05, Welch’s *t*-test) relative to the data obtained for the control strain.

For the identification of NrdA–mRNA interaction and evaluation of the destination of mRNA interacted with NrdA, we performed an experiment under the following conditions. The Tet-*S*-*nrdA* strain was precultivated in YES medium with Dox; next, mycelia were transferred to YES medium with or without Dox and further cultivated (Fig. 3C). The resultant mycelial cells cultivated with Dox were separated into two portions and used for RNA-seq analysis and RIP-seq analysis to identify the NrdA–mRNA interaction by comparing their RNA pools. In addition, the resultant mycelial cells cultivated without Dox were used for RNA-seq analysis to investigate the expression levels of NrdA-interacting mRNA in the presence or absence of Dox.

For the latter purpose, we evaluated the expression level of *S*-*nrdA* in the Tet-*S*-*nrdA* strain cultivated in the presence or absence of Dox. The Tet-*S*-*nrdA* strain was precultivated in YES medium with Dox for 16 h; next, mycelia were transferred to YES medium with or without Dox and further cultivated for 4, 6, 12, or 24 h. In addition, the *A. kawachii* control strain was precultivated in the YES medium for 16 h; subsequently, the mycelia were transferred to YES medium and further cultivated for 24 h. After the cultivations, total RNA was extracted and used to compare the expression levels of *S-nrdA* (Fig. 3D). The levels of *S*-*nrdA* in the Tet-*S*-*nrdA* strain cultivated in the medium with Dox was significantly higher (17.3–37.2-fold higher) than those of *nrdA* in the control strain, indicating that S-NrdA was overexpressed throughout the cultivation period. On the other hand, the expression level of *S-nrdA* in the Tet-*S*-*nrdA* strain cultivated in the medium without Dox was gradually decreased; however, it remained comparable to the expression level of *nrdA* in the control strain after cultivation for 12 h and 24 h. This result indicated that the *S*-*nrdA* transcript was not depleted even after cultivation without Dox for 24 h. Thus, we considered cultivation of the Tet-*S*-*nrdA* strain with and without Dox as *S*-*nrdA* overexpression and expression condition, respectively. To confirm this consideration, we additionally performed RNA-seq analysis of the control strain cultivated in YES medium without Dox (Fig. 3C, an experimental condition was surrounded by the dotted line), followed by clustering analysis of RNA-seq along with the RNA-seq and RIP-seq data obtained using the Tet-*S*-*nrdA* strain (Fig. 3E). The results indicated that the transcriptomic profile of the control strain is more similar to that of the Tet-*S*-*nrdA* strain cultivated without Dox than that of the Tet-*S*-*nrdA* strain cultivated with Dox. In addition, this is consistent with the data showing that the Tet-*nrdA* strain cultivated without Dox showed similar CAP compared with the control strain. In contrast, the Tet-*nrdA* strain cultivated with Dox exhibited significantly reduced CAP compared with the control strain (Fig. 1B).

### Analysis of NrdA-interacting transcripts and their destination

Based on the examination of experimental conditions, we cultivated the Tet-*S*-*nrdA* strain in YES medium with or without Dox as *S-nrdA* overexpression and expression condition, respectively, and the mycelia were subjected to RNA-seq and RIP-seq (Fig. 3C). A 4.4-fold higher amount of RNA was obtained from the mycelia of the Tet-*S-nrdA* strain cultivated with Dox by RIP using the anti-S-tag antibody compared with that obtained using the normal rabbit IgG (negative control) (Fig. 4A). This result indicated that the enrichment of NrdA-associated RNA was successful. To identify the NrdA–mRNA interaction, we compared the mRNA profiles of the Tet-*S-nrdA* strain cultivated with Dox obtained from the RNA-seq and RIP-seq analyses (Fig. 3C). We predicted NrdA– mRNA interaction by identifying the mRNA significantly enriched by RIP based on the following criteria: log_2_ fold change >0 and *q*-value <0.05 (see Data Set S1 in the supplemental material). This analysis identified 3,676 mRNA transcripts that represent 32% of the total 11,474 predicted coding sequences of *A. kawachii*. This is consistent with the estimation that Nrd1 and Nab3 directly regulate a variety of protein-coding genes, representing 20%–30% of the protein-coding transcripts of *S. cerevisiae* (5).

**FIG 4.**
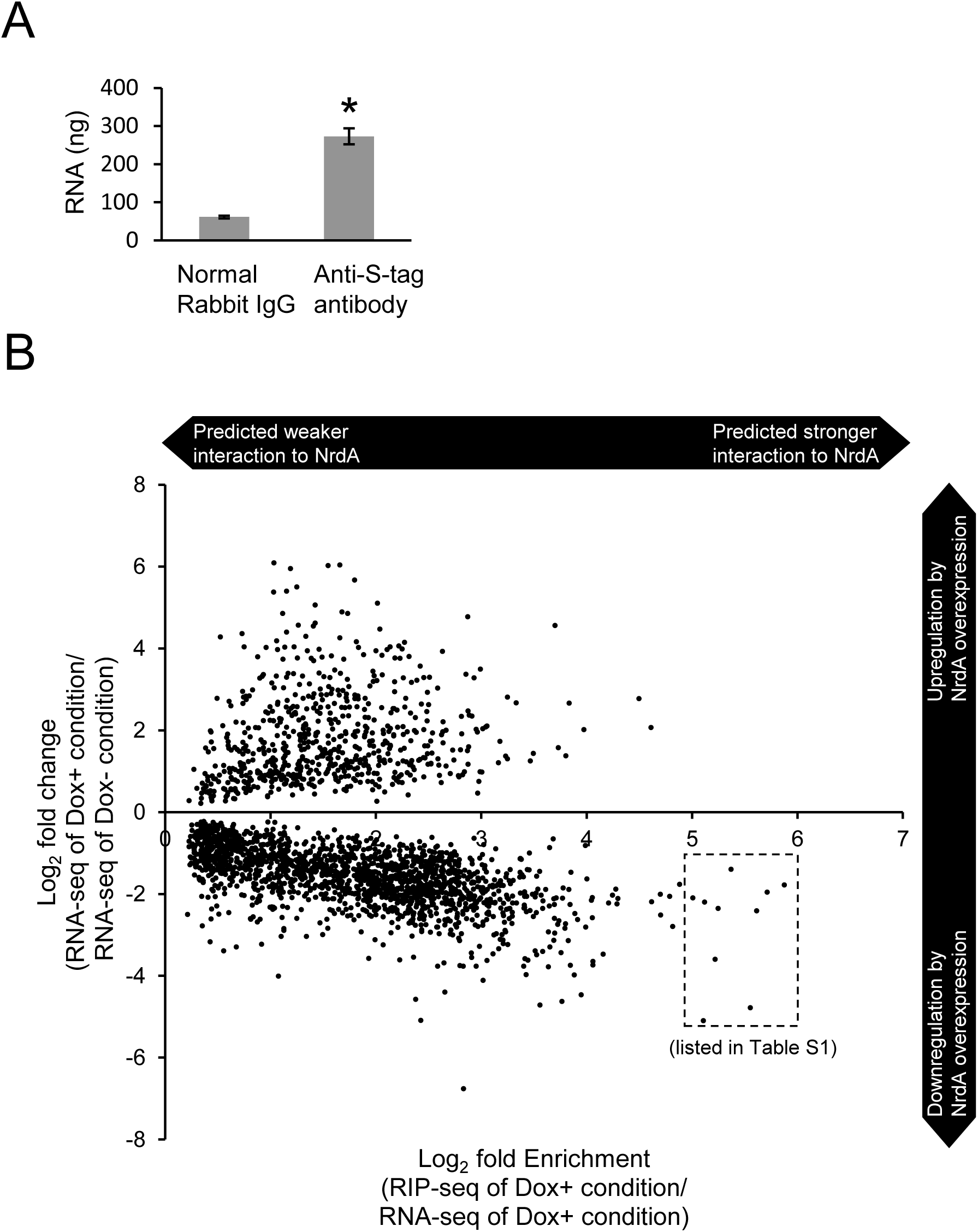
(A) The RNA obtained by immunoprecipitation from the Tet-*S*-*nrdA* strain cultivated in the presence of Dox. Anti-S-tag antibody and normal rabbit IgG (as a negative control) were used for the RNA immunoprecipitation. The obtained RNA was measured by NanoDrop-8000 (Thermo Fisher Scientific). The mean and standard deviation were determined from the results of 3 independent cultivations. *, statistically significant difference (*p* < 0.05, Welch’s *t*-test) relative to the negative control. (B) Change in gene expression of putative NrdA-interacting transcripts by the overexpression of NrdA. Predicted NrdA-interacting transcripts were plotted according to the log_2_ fold enrichment detected by the RIP analysis on the X-axis and the log_2_ fold change in gene expression observed after the overexpression of NrdA on the Y-axis. All the plotted data showed statistical significance (*q* < 0.05).

This study was initiated due to an interest in the *nrdA* genes localized on the syntenic region with *citA* and *yhmA* on the genome. However, the transcripts of *citA* (AKAW_06279) and *yhmA* (AKAW_06280) were not predicted to interact with NrdA (see Data Set S1 in the supplemental material). In addition, *citA* and *yhmA* are not essential genes in *A. kawachii* (11, data not shown), indicating that they are not responsible for the lethality caused by *nrdA* disruption.

We performed Gene Ontology (GO) enrichment analysis of mRNA transcripts, which were predicted to interact with NrdA. The results of this analysis identified metabolic process-, transport process-, and transcriptional regulation-related terms (Table 1). In addition, enriched terms included numerous secondary metabolic related terms, such as the secondary metabolic process (GO:0019748) and mycotoxin biosynthetic process (GO:0043386). It should be noted that *A. kawachii* does not produce mycotoxins, such as ochratoxin A and fumonisin B, and is considered safe for use in the food and beverage industry (22, 23).

**Table 1.** Biological process GO terms of potential NrdA-interacting mRNA identified by AsgGD GO Term Finder.

| GO ID | GO term | Number in gene set | Number in background | *P*-value |
| --- | --- | --- | --- | --- |
| GO:0019748 | secondary metabolic process | 160 | 310 | 1.23E-27 |
| GO:0055085 | transmembrane transport | 307 | 799 | 2.68E-24 |
| GO:0055114 | oxidation-reduction process | 339 | 984 | 1.49E-17 |
| GO:0044550 | secondary metabolite biosynthetic process | 115 | 243 | 2.21E-15 |
| GO:0008152 | metabolic process | 1226 | 4844 | 7.69E-09 |
| GO:0043386 | mycotoxin biosynthetic process | 56 | 113 | 1.80E-07 |
| GO:0043385 | mycotoxin metabolic process | 57 | 116 | 1.88E-07 |
| GO:0009404 | toxin metabolic process | 58 | 119 | 1.95E-07 |
| GO:0018958 | phenol-containing compound metabolic process | 43 | 77 | 2.01E-07 |
| GO:0009403 | toxin biosynthetic process | 56 | 115 | 4.28E-07 |
| GO:0051234 | establishment of localization | 378 | 1315 | 1.68E-06 |
| GO:0042537 | benzene-containing compound metabolic process | 31 | 50 | 2.45E-06 |
| GO:0018130 | heterocycle biosynthetic process | 237 | 766 | 3.76E-06 |
| GO:1901362 | organic cyclic compound biosynthetic process | 253 | 828 | 3.86E-06 |
| GO:0046189 | phenol-containing compound biosynthetic process | 35 | 61 | 4.15E-06 |
| GO:0006810 | transport | 369 | 1289 | 4.73E-06 |
| GO:1901376 | organic heteropentacyclic compound metabolic process | 44 | 86 | 5.18E-06 |
| GO:1901378 | organic heteropentacyclic compound biosynthetic process | 43 | 84 | 7.69E-06 |
| GO:0051179 | localization | 394 | 1402 | 1.47E-05 |
| GO:1903506 | regulation of nucleic acid-templated transcription | 216 | 697 | 1.76E-05 |
| GO:0006355 | regulation of transcription, DNA-templated | 216 | 697 | 1.76E-05 |
| GO:2001141 | regulation of RNA biosynthetic process | 216 | 698 | 2.01E-05 |
| GO:1900813 | monodictyphenone metabolic process | 23 | 34 | 3.07E-05 |
| GO:1900815 | monodictyphenone biosynthetic process | 23 | 34 | 3.07E-05 |
| GO:0051252 | regulation of RNA metabolic process | 219 | 714 | 3.57E-05 |
| GO:1900555 | emericalamide metabolic process | 22 | 32 | 4.03E-05 |
| GO:1900557 | emericalamide biosynthetic process | 22 | 32 | 4.03E-05 |
| GO:0050761 | depsipeptide metabolic process | 22 | 32 | 4.03E-05 |
| GO:0050763 | depsipeptide biosynthetic process | 22 | 32 | 4.03E-05 |
| GO:1901334 | lactone metabolic process | 24 | 37 | 5.21E-05 |
| GO:1901336 | lactone biosynthetic process | 24 | 37 | 5.21E-05 |
| GO:0042180 | cellular ketone metabolic process | 44 | 94 | 0.00016 |
| GO:0072330 | monocarboxylic acid biosynthetic process | 50 | 113 | 0.0002 |
| GO:1901503 | ether biosynthetic process | 22 | 34 | 0.00021 |
| GO:0030638 | polyketide metabolic process | 22 | 34 | 0.00021 |
| GO:0030639 | polyketide biosynthetic process | 21 | 32 | 0.00029 |
| GO:0005975 | carbohydrate metabolic process | 138 | 422 | 0.00032 |
| GO:0042181 | ketone biosynthetic process | 39 | 81 | 0.00034 |
| GO:1900582 | o-orsellinic acid metabolic process | 16 | 21 | 0.00038 |
| GO:1900584 | o-orsellinic acid biosynthetic process | 16 | 21 | 0.00038 |
| GO:1900552 | asperfuranone metabolic process | 20 | 30 | 0.00039 |
| GO:1900554 | asperfuranone biosynthetic process | 20 | 30 | 0.00039 |
| GO:1902644 | tertiary alcohol metabolic process | 20 | 30 | 0.00039 |
| GO:1902645 | tertiary alcohol biosynthetic process | 20 | 30 | 0.00039 |
| GO:0045460 | sterigmatocystin metabolic process | 36 | 73 | 0.00047 |
| GO:2000112 | regulation of cellular macromolecule biosynthetic process | 229 | 775 | 0.00048 |
| GO:0010556 | regulation of macromolecule biosynthetic process | 230 | 779 | 0.00049 |
| GO:0031326 | regulation of cellular biosynthetic process | 238 | 813 | 0.00063 |
| GO:0019438 | aromatic compound biosynthetic process | 211 | 707 | 0.00066 |
| GO:0045461 | sterigmatocystin biosynthetic process | 23 | 71 | 0.00071 |
| GO:0009889 | regulation of biosynthetic process | 240 | 822 | 0.00071 |
| GO:0034311 | diol metabolic process | 25 | 44 | 0.0011 |
| GO:0034312 | diol biosynthetic process | 25 | 44 | 0.0011 |
| GO:0019219 | regulation of nucleobase-containing compound metabolic process | 222 | 755 | 0.00112 |
| GO:1901615 | organic hydroxy compound metabolic process | 91 | 258 | 0.00116 |
| GO:1901617 | organic hydroxy compound biosynthetic process | 62 | 158 | 0.00127 |
| GO:0010468 | regulation of gene expression | 232 | 797 | 0.00145 |
| GO:0006351 | transcription, DNA-templated | 124 | 384 | 0.00284 |
| GO:0097659 | nucleic acid-templated transcription | 124 | 384 | 0.00284 |
| GO:0009820 | alkaloid metabolic process | 26 | 49 | 0.00387 |
| GO:0009821 | alkaloid biosynthetic process | 26 | 49 | 0.00387 |
| GO:0032774 | RNA biosynthetic process | 124 | 387 | 0.00443 |
| GO:0080090 | regulation of primary metabolic process | 253 | 899 | 0.00828 |
| GO:0018904 | ether metabolic process | 23 | 43 | 0.01271 |
| GO:0060255 | regulation of macromolecule metabolic process | 256 | 919 | 0.01666 |
| GO:0035834 | indole alkaloid metabolic process | 23 | 45 | 0.03425 |
| GO:0035835 | indole alkaloid biosynthetic process | 23 | 45 | 0.03425 |

Next, to examine the significance of NrdA–mRNA interaction, differential gene expression between the S-NrdA expression and overexpression conditions was compared using RNA-seq (see Data Set S1 in the supplemental material). A *q*-value <0.05 denoted statistically significant changes in gene expression. Based on this criterion, the predicted 3,676 NrdA-associated mRNAs included 1,768 downregulated genes and 674 upregulated genes by the overexpression of S-NrdA. These findings indicated that approximately half of NrdA-associated mRNAs were downregulated by the overexpression of S-NrdA. In addition, RIP-seq and RNA-seq plotting analyses showed that the proportion of downregulated genes was increased in the higher enriched mRNA transcript by RIP (predicted strong interaction of mRNA with S-NrdA) (Fig. 4B). For example, the higher enriched mRNA transcript (with log_2_ fold change >5) included 10 genes downregulated by the overexpression of S-NrdA (surrounded by dotted line in Fig. 4B). Based on the information obtained from domain search using the Pfam database, these may have beta lactamase (AKAW_10564), G-protein coupled receptor (AKAW_10304), aldehyde reductase (AKAW_01947), or tannanase (AKAW_10686 and AKAW_04699) functions; however, most of them are hypothetical proteins with unknown function (Table S1). In addition, *S. cerevisiae* and *Schiz. pombe* have homologs only for AKAW_01947 among these 10 genes. The physiological reason responsible for the interaction of the transcripts of these genes with NrdA remains unclear. Nevertheless, the overexpression of NrdA associated with transcriptional downregulation may be attributed to the negative effect exerted by the NNS complex pathway on the mRNA expression through the termination of transcription (3–5). Hence, the overexpression of S-NrdA may cause the downregulation of interacted RNA molecules by the transcription termination.

### NrdA regulates the production of secondary metabolites in *A. nidulans, A. fumigatus*, and *A. oryzae*

GO analysis indicated that the significant change observed in the transcriptional levels of secondary metabolism was related to genes induced by the overexpression of NrdA in *A. kawachii* (Table 1). This result was interesting because the relationships between the NNS complex pathway and the production of secondary metabolites have not been studied. However, the secondary metabolites of *A. kawachii* are currently largely unknown. Therefore, we tested the effect of overexpression of intrinsic NrdA on the production of secondary metabolites in *A. nidulans*, *A. fumigatus*, and *A. oryzae*.

*A. nidulans* OE-*nrdA* strain showed a slightly fluffy phenotype (Fig. 5A) with reduced number of conidia (Fig. 5B). To investigate the production of penicillin and sterigmatocystin, *A. nidulans* strains were precultivated in YES medium at 30°C for 16 h, transferred to minimal medium, and further cultivated at 30°C for 24, 48, or 72 h. The results of the halo assay indicated that the OE-*nrdA* strain showed significantly reduced penicillin production compared with the control strain (Fig. 5C). The sizes of the halos were significantly different using culture supernatant obtained after 24 and 48 h, whereas they were similar at 72 h. This evidence indicates that overexpression of NrdA suppressed the production of penicillin in the early cultivation period. In addition, the production of sterigmatocystin by OE-*nrdA* was 26% of that produced by the control strain after 24 h of cultivation (Fig. 5D, left side). This difference was reduced after 48 h of cultivation (Fig. 5D, right side). We confirmed that transcriptional levels of *nrdA* in OE-*nrdA* were 15-fold higher compared with those detected in the control strain at 12 h (Fig. 5E). The differences in the transcriptional levels of *nrdA* in OE-*nrdA* were not statistically significant at 24 and 48 h, indicating that *nrdA* is overexpressed in the early cultivation period of the OE-*nrdA* strain. Next, we investigated whether the reduced production of penicillin and sterigmatocystin was controlled at the transcriptional level. The transcriptional levels of *ipnA* (an isopenicillin-*N* synthase with a role in penicillin biosynthesis) (24), *aflR* (a transcriptional regulator involved in the production of sterigmatocystin) (25), and *stcU* (a putative versicolorin reductase involved in the biosynthesis of sterigmatocystin) (26, 27) were significantly reduced in the OE-*nrdA* strain at 12 and 24 h compared with the control strain. However, these levels became similar at 48 h. This is consistent with the data showing that the higher transcriptional level of *nrdA* and reduced productions of penicillin and sterigmatocystin were significant in the early cultivation period.

**FIG 5.**
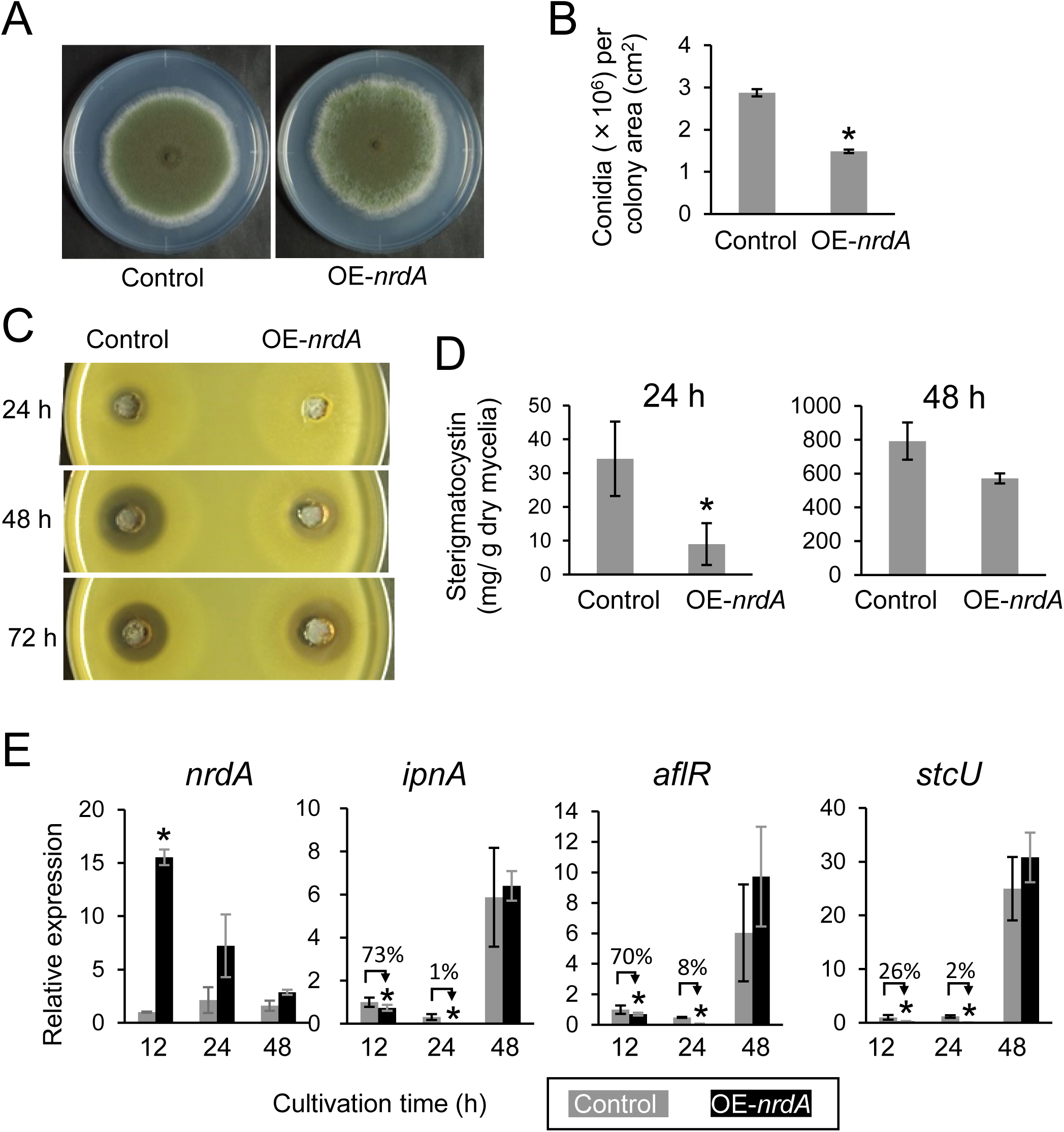
(A) Colony formation of the *A. nidulans* control and OE-*nrdA* strains. Conidia (1 × 10^4^) were inoculated onto minimal agar medium with biotin and cultured at 30°C for 7 days. (B) Conidiation of the control and OE-*nrdA* strains. (C) Penicillin bioassay of the *A. nidulans* control and OE-*nrdA* strains. (D) Production of sterigmatocystin by the control and OE-*nrdA* strains. (E) Transcriptional levels of *nrdA*, *ipnA*, *aflR*, and *stcU*. The control and OE-*nrdA* strains were precultured in YES medium for 16 h, transferred to minimal medium with biotin, and further cultured for 12, 24, and 48 h. The mean and standard deviation were determined from the results of 3 independent cultivations. Asterisks indicate significant differences (* *p* < 0.05, Welch’s *t*-test) versus the results obtained for the control strain.

The *A. fumigatus* OE-*nrdA* strain showed similar colony sizes (Fig. 6A) with reduced number of conidia (Fig. 6B). Next, we analyzed the levels of fumagillin, helvolic acid, and pyripyropene A in culture supernatant (Fig. 6C). *A. fumigatus* strains were cultivated in minimal liquid medium with fetal bovine serum (FBS) at 30°C for 24 h. The difference in the production of fumagillin between the control and OE-*nrdA* strains were not statistically significant. In contrast, the production of helvolic acid and pyripyropene by OE-*nrdA* was significantly reduced to 47% and 3% of that observed in the control strain, respectively. We confirmed that the OE-*nrdA* strain exhibited higher expression of *nrdA* compared with the control strain (Fig. 6D). Subsequently, we investigated the gene expression of *fumR* (a putative C6 type transcription factor-encoding gene involved in fumagillin biosynthesis) (28), *helA* (an oxidosqualene cyclase-encoding gene involved in the production of helvolic acid) (29), and *pyr2* (a polyketide synthase-encoding gene involved in the biosynthesis of pyripyropene A) (30). Although the transcriptional levels of *fumR* in the control and OE-*nrdA* strains were similar, those of *helA* and *pyr2* were lower in the OE-*nrdA* strain (Fig. 6D). This was consistent with the results demonstrating that the production of helvolic acid and pyripyropene A was significantly suppressed by the overexpression of NrdA.

**FIG 6.**
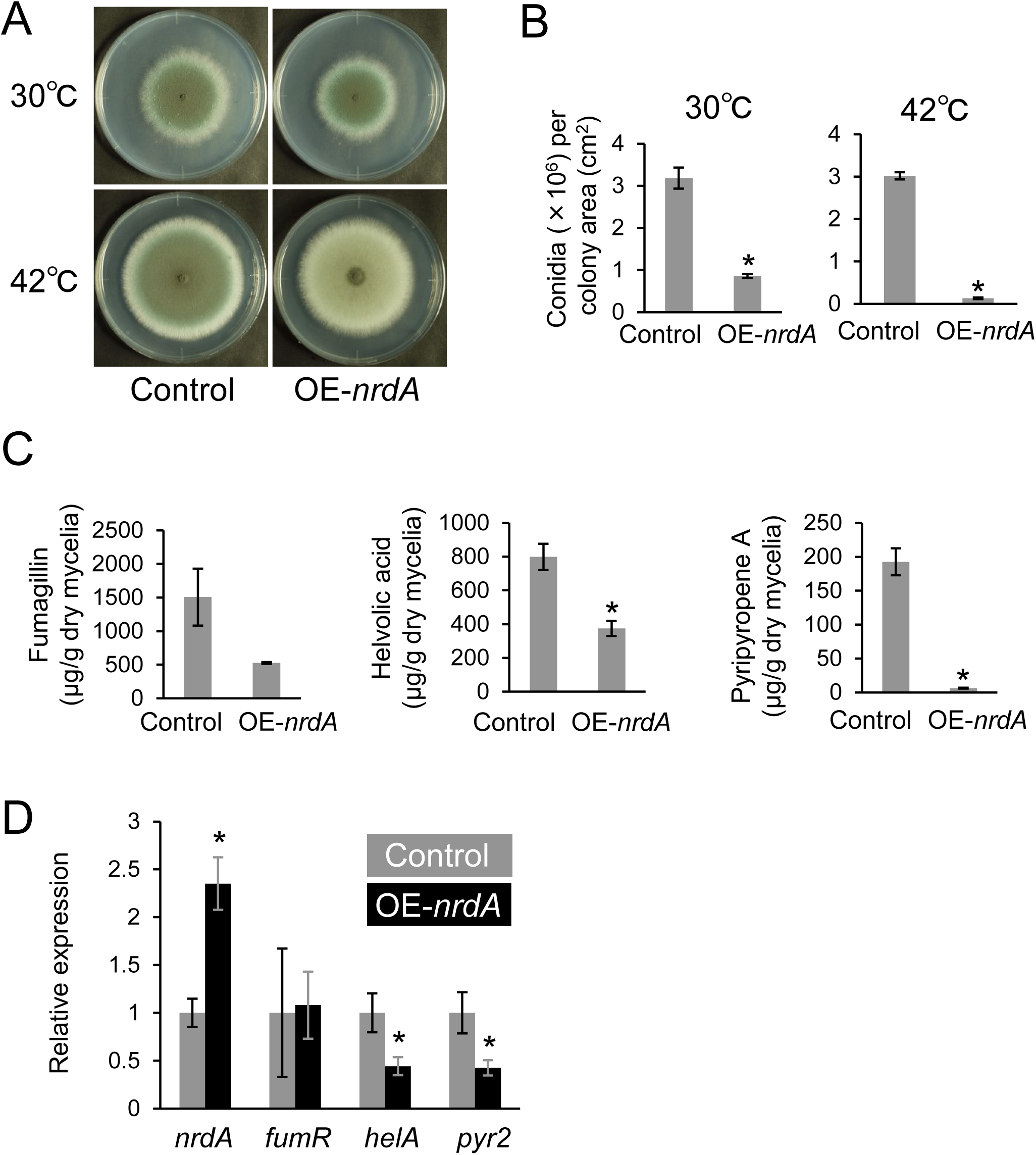
(A) Colony formation of the *A. fumigatus* control and OE-*nrdA* strains. Conidia (1 × 10^4^) were inoculated onto minimal agar medium and cultured at 30°C or 42°C for 5 days. (B) Conidiation of the control and OE-*nrdA* strains. (C) Production of fumagillin, helvolic acid, and pyripyropene A by the control and OE-*nrdA* strains. Conidia (2 × 10^7^ cells) were inoculated in minimal liquid medium with 5% FBS and cultivated with shaking (163 rpm) at 30°C for 24 h. (D) Transcriptional levels of *nrdA*, *fumR*, *helA*, and *pyr2*. The strains were cultivated in minimal liquid medium with 5% FBS for 12 h. The mean and standard deviation were determined from the results of 3 independent cultivations. Asterisks indicate significant differences (* *p* < 0.05, Welch’s *t*-test) versus the results obtained for the control strain.

*A. oryzae* control and OE-*nrdA* strains were cultivated in medium containing ferric ion (31), whose color turns red if it is chelated by kojic acid (Fig. 7A). The OE-*nrdA* strain showed a reduced number of conidia (Fig. 7B). In addition, the number of sclerotia increased in the OE-*nrdA* strain compared with the control strain (Fig. 7A). The bottom side of the colony of the OE-*nrdA* strain exhibited a red color, indicating that this strain produces higher amounts of kojic acid compared with the control strain. To investigate the transcriptional levels of genes involved in the production of kojic acid, the strains were cultivated in kojic acid production liquid medium at 30°C for 5 days. Furthermore, we confirmed a higher transcriptional level of *nrdA* in the OE-*nrdA* strain compared with that detected in the control strain (Fig. 7C). In addition, the gene expression of *kojR*, *kojA*, and *kojT*, which are involved in the production of kojic acid (31), also exhibited higher levels compared with those measured in the control strain. To investigate the production of penicillin, strains were precultured in YES medium, transferred to minimal medium, and further cultivated for 24 or 48 h. The results of the halo assay using the culture supernatant at 48 h revealed significantly reduced production of penicillin by the OE-*nrdA* strain (Fig. 7D). In addition, the transcriptional analysis confirmed that the OE-*nrdA* strain showed higher and similar expression levels of *nrdA* compared with the control strain at 12 and 24 h, respectively (Fig. 7E). In addition, the OE-*nrdA* strain showed higher and lower transcriptional levels of *ipnA* compared with the control strain at 12 and 24 h, respectively. The significantly lower transcriptional level of *ipnA* is consistent with the data indicating that penicillin production was suppressed by the overexpression of *nrdA* at 48 h.

**FIG 7.**
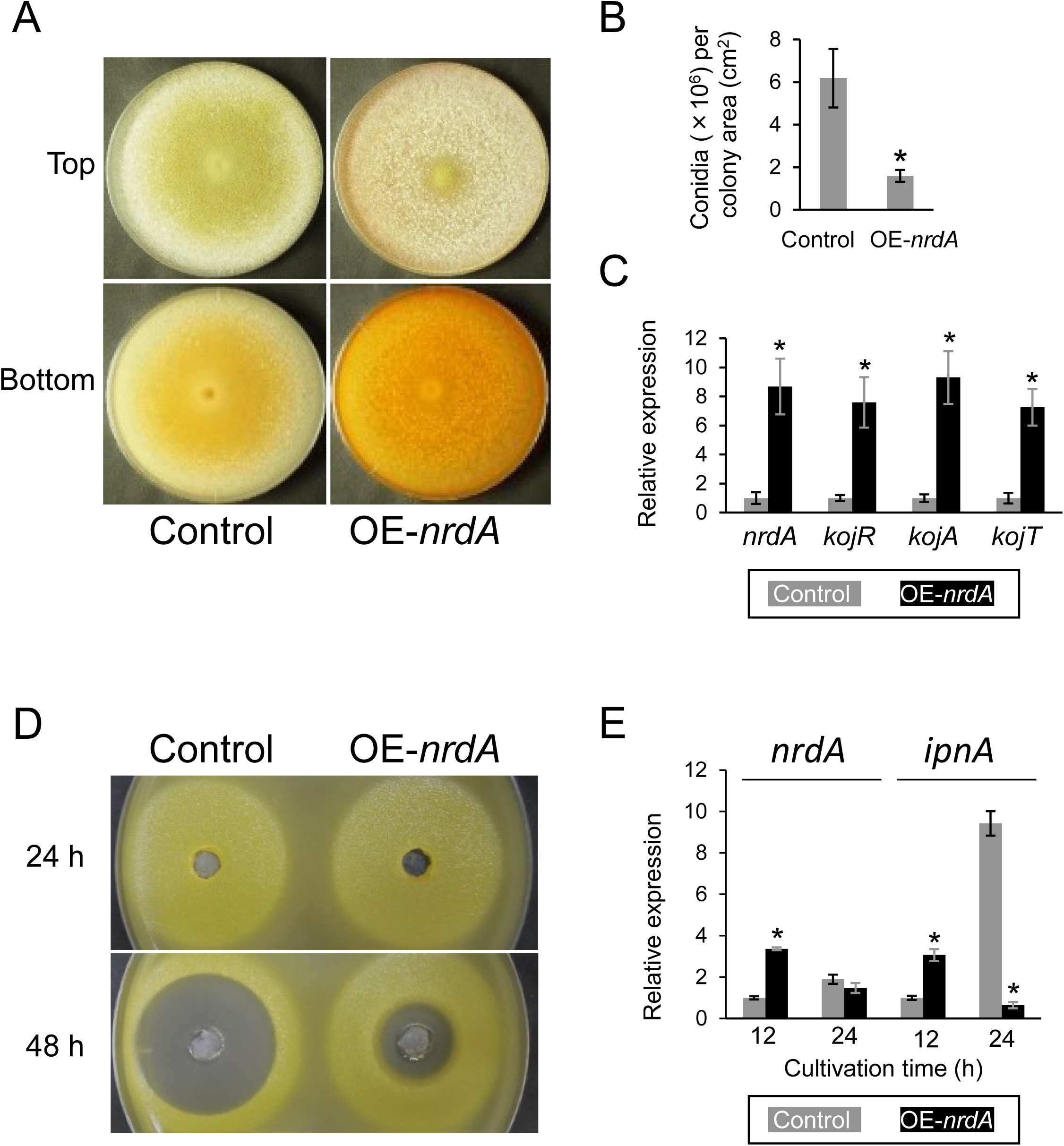
(A) Colony formation of the *A. oryzae* control and OE-*nrdA* strains. Conidia (1 × 10^4^) were inoculated onto kojic acid production agar medium and cultured at 30°C for 5 days. (B) Conidiation of the control and OE-*nrdA* strains. (C) Transcriptional levels of *nrdA*, *kojR*, *kojA*, and *kojT*. Conidia (2 × 10^7^ cells) were inoculated in kojic acid production liquid medium and cultivated with shaking (163 rpm) at 30°C for 5 days. (D) Penicillin bioassay of the *A. oryzae* control and OE-*nrdA* strains. (E) Transcriptional levels of *nrdA* and *ipnA*. The mean and standard deviation were determined from the results of 3 independent cultivations. Asterisks indicate significant differences (* *p* < 0.05, Welch’s *t*-test) versus the results obtained for the control strain.

The overexpression of *nrdA* significantly affected the production of secondary metabolites in *A. nidulans*, *A. fumigatus*, and *A. oryzae*. A hypothesis is that NrdA may be directly involved in the silencing of secondary metabolite gene clusters via a NNS-related transcription termination system because the overexpression of *nrdA* suppressed the gene expression levels of *ipnA*, *aflR*, and *stcU* in *A. nidulans* (Fig. 5E), *helA* and *pyr2* in *A. fumigatus* (Fig. 6D), and *ipnA* in *A. oryzae* (Fig. 7E). Another hypothesis is that these genes may be indirectly regulated via transcriptional factors and/or histone modification and chromatin remodeling, which often regulate the production of secondary metabolites in fungi (32). The transcriptional factor BrlA regulates the production of fumagillin, helvolic acid, and pyripyropene A in *A. fumigatus* (33). The production of sterigmatocystin and penicillin is regulated by the putative methyltransferase LaeA in *A. nidulans* (34). The production of penicillin and kojic acid is regulated by histone deacetylase Hst4 and LaeA (35, 36). In *S. cerevisiae*, Nrd1 and exosome-dependent RNA degradation leads to gene silencing by histone modification. The euchromatin mark, such as histone H3 trimethylated at lysine 4 (H3K4me3) and histone H3 acetylated at lysine 9 (H3K9ac) at non-transcribed spacer locus of ribosomal DNA, a heterochromatin region is enriched by the depletion of Nrd1 (37). Recently, it was also reported that heterochromatin assembly occurs by the pausing of RNA polymerase II promoted by Seb1 in *Schiz. pombe* (38). Thus, overexpression of NrdA might contribute to the formation of heterochromatin at the secondary metabolite gene cluster regions in *Aspergillus* spp.

We could not determine the reason for the co-localization of *nrdA* with *citA* and *yhmA* in subdivision Pezizomycotina; however, NrdA plays a significant role in global gene regulation and is involved in the production of secondary metabolites in *Aspergillus* species. CAP is regulated by the regulator of secondary metabolism LaeA in *Aspergillus niger*, *Aspergillus carbonarius*, and *A. kawachii* (39–41). Therefore, NrdA may be linked to CAP via the regulation of secondary metabolism. Although the NNS-associated RNA surveillance system has been extensively investigated in the yeasts *S. cerevisiae* and *Schiz. pombe*, the present findings may enhance the understanding of the NNS-associated production of secondary metabolites in *Aspergillus* spp., including industrially and clinically important fungi.

## MATERIALS AND METHODS

### Strains and culture conditions

In this study, *A. kawachii* SO2 (42), *A. nidulans* A26 (Fungal Genetics Stock Center [FGSC; Manhattan, KS]), *A. fumigatus* A1151 (FGSC), and *A. oryzae* Δ*ligD* plus pGNA (43, 44) were used as parental strains (Table 2). Control strains were defined to show an identical auxotrophic background for comparison with the respective *nrdA* overexpression strains.

**Table 2.**
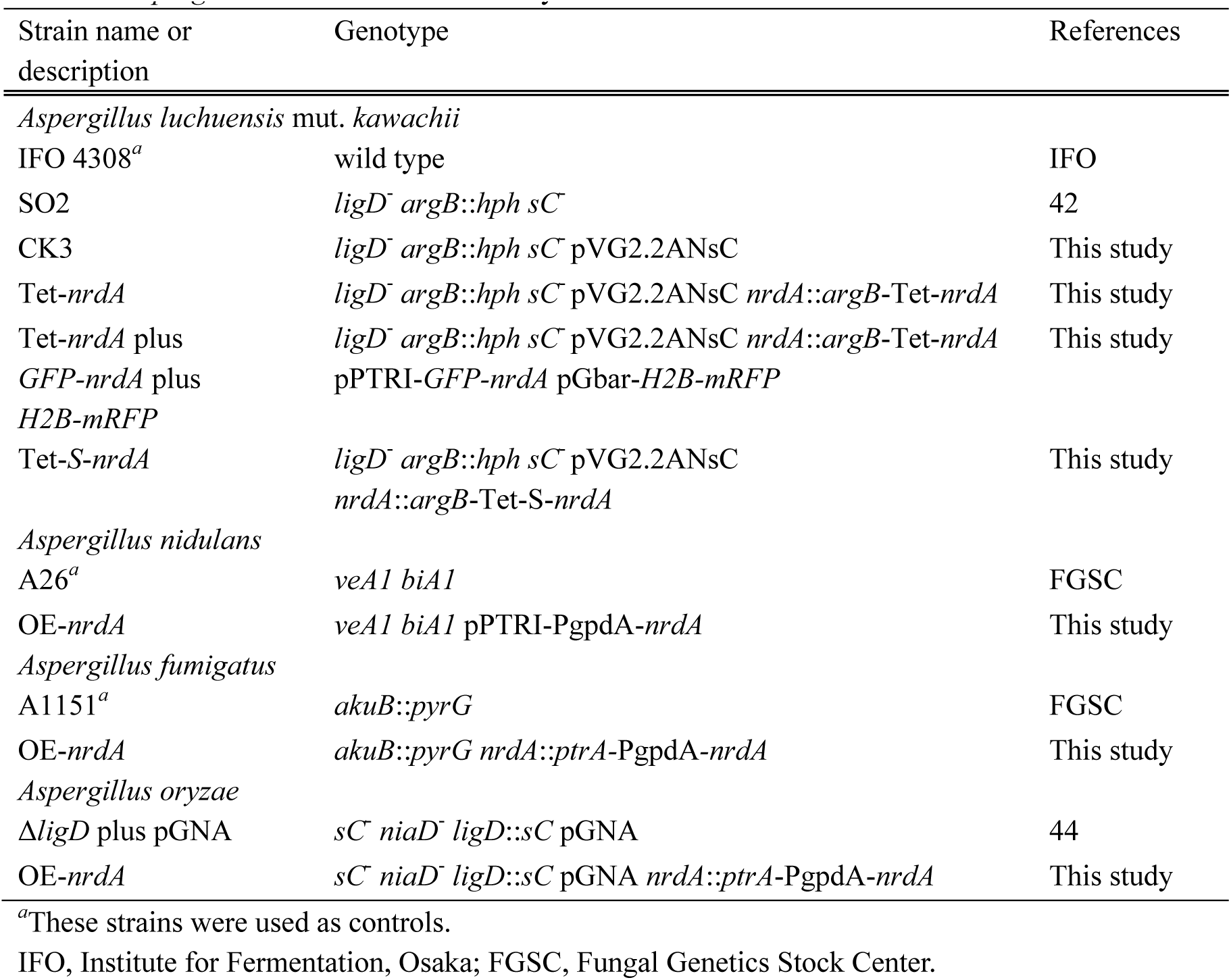
Aspergillus strains used in this study

Strains were cultivated in minimal medium (45; FGSC [http://www.fgsc.net/methods/anidmed.html]) with or without 0.211% (wt/vol) arginine and/or 0.15% (wt/vol) methionine, and/or 0.1 µg/ml pyrimethamine. CAP medium (11), kojic acid production medium with 1 mM ferric ion (31), or YES (15% sucrose [wt/vol], 2% yeast extract [wt/vol]) medium were used appropriately for the fungal growth experiments. Minimal and CAP media were adjusted to the required pH with NaOH and HCl, respectively. For the cultivation of *A. nidulans* strains, minimal medium was supplemented with 5 µg/ml biotin.

### Construction of *A. kawachii* Tet-*nrdA* and Tet-*S-nrdA* strains

To achieve Dox-inducible conditional expression of the *nrdA* gene, 2 kb of the 5’-end of *nrdA*, 2.2 kb of *argB*, 0.8 kb of *cgrA* terminator and Tet-On promoter, and 1.8 kb of a part of the ORF region of *nrdA* were constructed by recombinant PCR using the primer pairs AKtet-nrdA-FC/AKtet-nrdA-R1, AKtet-nrdA-F2/AKtet-nrdA-R2, AKtet-nrdA-F3/AKtet-nrdA-R3, and AKtet-nrdA-F4/AKtet-nrdA-RC, respectively (see Table S2 in the supplemental material). For amplification of the *argB* gene and Tet-On promoter, *A. nidulans* A26 genomic DNA and plasmid pVG2.2ANsC were used as template DNA, respectively (11, 46). The resultant DNA fragment was amplified with the primers AKtet-nrdA-F1 and AKtet-nrdA-R4 and used to transform the *A. kawachii* strain CK3, which carried pVG2.2ANsC, yielding the Tet-*nrdA* strain. Transformants were selected on minimal agar medium without arginine. Introduction of this cassette into the target locus was confirmed with PCR using the primer pairs AKtet-nrdA-FC and AKtet-nrdA-RC (see Fig. S3 in the supplemental material).

To confirm the functional expression of S-tagged NrdA, we constructed a S-NrdA expression strain (data not shown). The 2 kb of the 5’-end of *nrdA*, 1.8 kb of *ptrA*, and 1.8 kb of a part of the ORF region of S-*nrdA* were constructed by recombinant PCR using the primer pairs Aktet-nrdA-FC/AkS-nrdA-ptrA-R1, AkS-nrdA-ptrA-F2/AkS-nrdA-ptrA-R2, and AkS-nrdA-ptrA-F3/AKtet-nrdA-RC, respectively (see Table S2 in the supplemental material). For amplification of the *ptrA* gene, pPTR I (Takara bio, Shiga, Japan) (47) was used as template DNA. The resultant DNA fragment was amplified with the primers AKtet-nrdA-F1 and AKtet-nrdA-R4 and used to transform the *A. kawachii* strain SO2, yielding the *S-nrdA* strain. Transformants were selected on minimal agar medium with pyrithiamine.

To achieve Dox-inducible conditional expression of the *S*-*nrdA* gene, 2 kb of the 5’-end of *nrdA*, 2.2 kb of *argB*, 0.8 kb of *cgrA* terminator and Tet-On promoter, and 2 kb of a part of ORF region of *nrdA* were constructed by recombinant PCR using the primer pairs AKtet-nrdA-FC/AKtet-S-nrdA-R3, and AKtet-S-nrdA-F4/AKtet-nrdA-RC, respectively (see Table S2 in the supplemental material). For amplification of the 5’-end *nrdA*-*argB*-Tet-On promoter and S-*nrdA*, the genomic DNA of strain Tet-*nrdA* and strain S-*nrdA* were used as template DNA, respectively. The resultant DNA fragment was amplified with the primers AKtet-nrdA-F1 and AKtet-nrdA-R4 and used to transform the *A. kawachii* strain CK3, yielding the Tet-*S*-*nrdA* strain. Transformants were selected on minimal agar medium without arginine. Introduction of this cassette into the target locus was confirmed with PCR using the primer pairs AKtet-nrdA-FC and AKtet-nrdA-RC (see Fig. S3 in the supplemental material). The expression of S-tagged NrdA was confirmed by purification using S-protein agarose (Merck Millipore, Darmstadt, Germany) as previously described (11, 48). The purified protein was subjected to sodium dodecyl sulfate-polyacrylamide gel electrophoresis and immunoblotting analysis using an anti-S-tag antibody (Medical and Biological Laboratories, Nagoya, Japan) (Fig. 3B).

### Construction of an *A. kawachii* strain expressing GFP-NrdA and H2B-mRFP

The plasmid pPTR I (Takara Bio), which carries the *ptrA* gene, was used to construct the expression vector for GFP-NrdA. The 5’-end of *nrdA* (until start codon) and *gfp* and *nrdA* ORF (without start codon) were amplified by PCR using the primer pairs pPTRI-gfp-nrdA-F1/pPTRI-gfp-nrdA-R1, pPTRI-gfp-nrdA-F2/pPTRI-gfp-nrdA-R2, and pPTRI-gfp-nrdA-F3/pPTRI-gfp-nrdA-R3, respectively. For the amplification of *gfp*, plasmid pFNO3 (49) was used as template DNA. The resultant DNA fragment was amplified with the primers pPTRI-gfp-nrdA-inf-F/pPTRI-gfp-nrdA-inf-R (see Table S2 in the supplemental material). The amplified fragment was cloned into the SmaI site of pPTR I using an In-Fusion HD cloning kit (Takara Bio). The pPTR I-*GFP*-*nrdA* plasmid was used to transform the *A. kawachii* strain Tet-*nrdA*, yielding the GFP-NrdA strain. Transformants were selected on minimal agar medium with pyrithiamine. After fluorescence microscopy testing, the pGbar-H2B-mRFP plasmid (50) was used to transform the *A. kawachii* strain GFP-NrdA, yielding the GFP-NrdA H2B-mRFP strain. Transformants were selected on minimal agar medium with glufosinate extracted from the herbicide Basta (Bayer Crop Science, Bayer Japan, Tokyo, Japan).

### Fluorescence microscopy

A strain expressing GFP-NrdA and H2B-mRFP was cultured in minimal medium supplemented with arginine. After cultivation in minimal medium supplemented with arginine for 12 h, the mycelia were observed under a DMI6000B inverted-type fluorescent microscope (Leica Microsystems, Wetzlar, Germany). Image contrast was adjusted using the LAS AF Lite software, version 2.3.0, build 5131 (Leica Microsystems).

### Measurement of citric acid

To measure the levels of extracellular citric acid, conidia (2 × 10^7^ cells) of the *A. kawachii* control strain were inoculated into 100 ml of YES medium and precultivated with shaking (180 rpm) at 30°C for 16 h. Subsequently, they were transferred to 50 ml of CAP medium with arginine and further cultivated with shaking (163 rpm) at 30°C for 48 h. Prior to their transfer to CAP medium, the mycelia were washed using fresh CAP medium. The Tet-*nrdA* strain was precultured in YES medium with 1 µg/ml Dox and transferred to CAP medium with or without Dox. The culture supernatant was filtered through a PTFE filter (pore size: 0.2 μm) (Toyo Roshi Kaisha, Tokyo, Japan) and used as the extracellular fraction.

The concentration of citric acid was determined using a Prominence HPLC system (Shimadzu, Kyoto, Japan) equipped with a CDD-10AVP conductivity detector (Shimadzu). The organic acids were separated with tandem Shimadzu Shim-pack SCR-102H columns (300 × 8 mm I.D.; Shimadzu) at 50°C using 4 mM *p*-toluenesulfonic acid monohydrate as the mobile phase at a flow rate of 0.8 ml/min. The flow rate of the post-column reaction solution (4 mM *p*-toluenesulfonic acid monohydrate, 16 mM bis-Tris, and 80 µM ethylenediaminetetraacetic acid [EDTA]) was 0.8 ml/min.

### RNA preparation

To identify the transcriptome of *A. kawachii* control and Tet-*S-nrdA* strains, conidia (2 × 10^7^ cells) were inoculated into 100 ml of YES medium with 1 µg/ml Dox (for Tet-*S-nrdA* strain) and precultured for 14 h at 30°C. Subsequently, they were transferred to 50 ml of YES medium with or without Dox (for Tet-*S-nrdA* strain) and further cultivated with shaking (163 rpm) at 30°C for 24 h. Prior to their transfer to YES medium with or without Dox, the mycelia were washed with fresh YES medium. Following incubation, the mycelia were collected and ground to a powder in the presence of liquid nitrogen. Total RNA was extracted using RNAiso Plus (Takara Bio) according to the instructions provided by the manufacturer and quantified using a NanoDrop-8000 (Thermo Fisher Scientific).

### RIP assay

To identify NrdA interacting RNA, *A. kawachii* Tet-*S*-*nrdA* strain was cultivated using YES medium with Dox under the condition described for the RNA preparation. The mycelia were collected and ground to a powder in the presence of liquid nitrogen. The powdered mycelia (1 g wet weight) were dissolved in 13 ml of ice-cold nuclear extraction buffer (25 mM N-2-hydroxyethylpiperazine-N’-2-ethane sulfonic acid [HEPES; pH 6.8], 1 M sorbitol, 250 μg/ml phenylmethylsulfonyl fluoride [PMSF], cOmplete [EDTA-free protease inhibitor cocktail; Roche, Basel, Switzerland]) and vigorously mixed using a vortexer. Cell debris was removed by filtration using Miracloth, and the solution was centrifuged at 10,000 × *g* at 4°C for 15 min. The supernatant was removed and the pellet was dissolved in 500 µl of ice-cold nuclear solubilization buffer (25 mM HEPES [pH 6.8], 1 M sorbitol, 0.5% NP-40, 250 μg/ml PMSF, cOmplete [EDTA-free protease inhibitor cocktail; Roche]). Following incubation for 30 min at 4°C, the debris was removed by centrifugation at 2,000 × *g* at 4°C for 15 min. Twenty-five µL of Protein A Sepharose beads (50% slurry contained phosphate-buffered saline buffer) (GE healthcare, Chicago, IL) was added to the resultant supernatant, and the resulting mixture was gently mixed for 1 h at 4°C using a rotator. The mixture was separated by centrifugation at 2,000 × *g* at 4°C for 1 min, and 25 µl of anti-S-tag antibody (Medical and Biological Laboratories) or normal rabbit IgG (Medical and Biological Laboratories) cross-linked protein A Sepharose was added to the supernatant. Subsequently, these mixtures were gently mixed for 3 h at 4°C using a rotator. The RNA interacted with NrdA was purified using RiboCluster Profiler (Medical and Biological Laboratories) according to the instructions provided by the manufacturer and quantified using a NanoDrop-8000 (Thermo Fisher Scientific).

Library preparation, sequencing, and data analysis for RNA-seq and RIP-seq were performed by Kabushiki Kaisha DNAFORM (Yokohama, Japan). All RNA samples were treated with RiboZero (Human/Mouse/Rat) (Illumina, San Diego, CA) for the depletion of ribosomal RNA. All RNA-seq and RIP-seq experiments were performed thrice with RNA samples obtained from independently prepared mycelia using the NextSeq 500 system (Illumina) and mapped to the *A. kawachii* IFO 4308 genome (51) using a pipeline of trimming (trim_galore [http://www.bioinformatics.babraham.ac.uk/projects/trim_galore/], trimmomatic [52], cutadapt [53]), mapping (STAR) (54), counting gene levels (featureCount) (55), differential expression analysis (DEseq2) (56), and clustering analysis (MBCluster.Seq) (57). A gene set of *A. kawachii* were converted to a homologous gene set of *A. niger* to perform the GO term enrichment analysis using the GO Term Finder of the *Aspergillus* genome database (AspGD, http://www.aspgd.org/) (58). This species was used because *A. niger* is closely related to *A. kawachii* (9, 59, 60).

### Construction of the *A. nidulans nrdA* overexpression strain

The plasmid pPTR I (Takara Bio) was used to construct the overexpression vector for the *A. nidulans nrdA* gene (Locus tag, AN8276). The *A. nidulans gpdA* promoter and *nrdA* were amplified by PCR using the primer pairs pPTRI-PgpdA-nrdA-inf-F1/pPTRI-PgpdA-nrdA-inf-R1 and pPTRI-PgpdA-nrdA-inf-F2/pPTRI-PgpdA-nrdA-inf-R2 (see Table S2 in the supplemental material). The amplified fragments were cloned into the SmaI site of pPTR I using an In-Fusion HD cloning kit (Takara Bio). Transformants were selected on minimal agar medium with pyrithiamine.

### Construction of the *A. fumigatus nrdA* overexpression strain

For the overexpression analysis of *nrdA* in *A. fumigatus* (Locus tag, AFUB_052760), a gene replacement cassette encompassing a homology arm at the 5’ end of *nrdA*, *ptrA* selection marker, *A. nidulans gpdA* promoter, and homology arm at the *nrdA* locus was constructed through recombinant PCR using the primer pairs AfPgpdA-nrdA-FC/AfPgpdA-nrdA-R1, AfPgpdA-nrdA-F2/PgpdA-nrdA-R2, AfPgpdA-nrdA-F3/PgpdA-nrdA-R3, and AfPgpdA-nrdA-F4/AfPgpdA-nrdA-RC, respectively (see Table S2 in the supplemental material). For the amplification of DNA fragments, *A. fumigatus* FGSC A1151 genomic DNA, *A. nidulans* A26 genomic DNA, or plasmid pPTR I (Takara Bio) were used as template DNA. The resultant DNA fragments amplified with primers AfPgpdA-nrdA-F1/AfPgpdA-nrdA-R4 were used to transform the *A. fumigatus* FGSC A1151, yielding the *A. fumigatus* OE-*nrdA* strain. Transformants were selected on minimal agar medium supplemented with 0.1 µg/ml pyrithiamine. Introduction of the *ptrA* and *A. nidulans gpdA* promoter into the target locus was confirmed by PCR using primers AfPgpdA-nrdA-FC/AfPgpdA-nrdA-RC (see Fig. S4 in the supplemental material).

### Construction of the *A. oryzae nrdA* overexpression strain

For the overexpression analysis of *nrdA* in *A. oryzae* (Locus tag, AO090102000629), a gene replacement cassette encompassing a homology arm at the 5’ end of *nrdA*, *ptrA* selection marker, *A. nidulans gpdA* promoter, and homology arm at the *nrdA* locus of *A. oryzae* was constructed with recombinant PCR using the primer pairs AoPgpdA-nrdA-FC/AoPgpdA-nrdA-R1, AoPgpdA-nrdA-F2/PgpdA-nrdA-R2, AoPgpdA-nrdA-F3/PgpdA-nrdA-R3, and AoPgpdA-nrdA-F4/AoPgpdA-nrdA-RC, respectively (see Table S2 in the supplemental material). For the amplification of DNA fragments, *A. oryzae* RIB40 wild-type genomic DNA, *A. nidulans* A26 genomic DNA, and plasmid pPTR I (Takara Bio) were used as template DNA. These resultant DNA fragments amplified with primers AoPgpdA-nrdA-F1/AoPgpdA-nrdA-R4 were used to transform the *A. oryzae* strain Δ*ligD* plus pGNA (43, 44), yielding the *A. oryzae* strain OE-*nrdA*. Transformants were selected on minimal agar medium supplemented with 0.1 µg/ml pyrithiamine. Introduction of the *ptrA* and *A. nidulans gpdA* promoter into the target locus was confirmed by PCR using primers AoPgpdA-nrdA-FC/AoPgpdA-nrdA-RC (see Fig. S5 in the supplemental material).

### Analysis of secondary metabolites by liquid chromatography-mass spectrometry

Conidia (2 × 10^7^ cells) were inoculated into 100 ml of YES medium and precultivated with shaking (180 rpm) at 30°C for 16 h. Subsequently, they were transferred to 50 ml of minimal medium supplemented with biotin (for *A. nidulans* strains) or with 5% (vol/vol) FBS (for *A. fumigatus* strains) and further cultivated with shaking (163 rpm) at 30°C for 24 or 48 h. The mycelia were collected and used to measure the weight of freeze-dried mycelia. Culture supernatant (5 ml) was collected and mixed with an equal volume of ethyl acetate for *A. nidulans* or chloroform for *A. fumigatus*. The mixture was centrifuged at 10,000 × *g* at 4°C for 15 min. The ethyl acetate fraction was collected, evaporated by N_2_ gas spraying, and resuspended in 100 µl of acetonitrile. The concentration of sterigmatocystin, fumagillin, helvolic acid, and pyripyropene A was determined using a Prominence ultra-high-performance liquid chromatograph system (Shimadzu) equipped with a 3200 QTRAP system (AB SCIEX, Framingham, MA) in the multiple reaction monitoring mode. The secondary metabolites were separated with an ACQUITY UPLC CSH C18 Column (1.7 µm, 2.1 mm × 50 mm) (Waters, Milford, MA) at 40°C using 0.05% formic acid in acetonitrile as the organic phase and 0.05% formic acid in water as the aqueous phase at a flow rate of 0.2 ml/min. The solvent gradient started at 20% organic for 2 min, followed by a linear increase to 60% organic over 10 min, a linear increase to 100% organic over 1 min, and final maintenance at 100% organic for 20 min.

### Penicillin bioassay

Conidia (2 × 10^7^ cells) of the *A. nidulans* control and OE-*nrdA* strains and *A. oryzae* control and OE-*nrdA* strains were inoculated into 100 ml of YES medium and precultivated with shaking (180 rpm) at 30°C for 16 h. Subsequently, they were transferred to 50 ml of minimal medium supplemented with biotin and further cultivated with shaking (163 rpm) at 30°C for 24, 48, or 72 h. Culture supernatant (5 ml) was evaporated by freeze drying and resuspended in 500 µl of water. Penicillin bioassay was performed using *Kocuria rhizophila* (NBRC 12708) as previously reported (34).

### Kojic acid assay

Conidia (1 × 10^4^ cells) of *A. oryzae* strains were inoculated onto kojic acid production agar medium with ferric ion (chelation by kojic acid changes the color of the medium to red) (31). After cultivation at 30°C for 5 days, the color of the medium was observed to evaluate the production of kojic acid.

### Transcriptional analysis

To evaluate the expression level of *S*-*nrdA* in the *A. kawachii* Tet-*S*-*nrdA* strain, the Tet-*S*-*nrdA* strain was precultivated in YES medium with Dox for 16 h. Next, the mycelia were transferred to YES medium with or without Dox and further cultivated for 4, 6, 12, or 24 h with shaking (163 rpm) at 30°C. In addition, the *A. kawachii* control strain was precultivated in the YES medium for 16 h. The mycelia were transferred to YES medium and further cultivated for 24 h.

To evaluate the gene expression level involved in the production of secondary metabolites, *A. nidulans*, *A. fumigatus*, and *A. oryzae* strains were cultivated under the condition described for the analysis of secondary metabolites. For the analysis of transcripts related to the production of kojic acid in *A. oryzae*, conidia (2 × 10^7^ cells) were inoculated onto kojic acid production medium with ferric ion (31) and incubated with shaking (163 rpm) at 30°C for 5 days. The mycelia were collected from the liquid culture using a gauze and ground to a powder in the presence of liquid nitrogen.

Total RNA was extracted using RNAiso Plus (Takara Bio) according to the instructions provided by the manufacturer and quantified using a NanoDrop-8000 (Thermo Fisher Scientific). cDNA was synthesized from total RNA using a PrimeScript Perfect real-time reagent kit (Takara Bio) according to the instructions provided by the manufacturer. Real-time reverse transcription-PCR was performed using a Thermal Cycler Dice real-time system MRQ (Takara Bio) with TB Green Premix Ex Taq II (Tli RNaseH Plus) (Takara Bio). The following primer sets were used: ANnrdA-RT-F and ANnrdA-RT-R for *A. nidulans nrdA*, ANipnA-RT-F and ANipnA-RT-R for *A. nidulans ipnA*, ANaflR-RT-F and ANaflR-RT-R for *A. nidulans aflR*, ANstcU-RT-F and ANstcU-RT-R for *A. nidulans stcU*, ANacnA-RT-F and ANacnA-RT-R for *A. nidulans acnA*, AFnrdA-RT-F and AFnrdA-RT-R for *A. fumigatus nrdA*, AFfumR-RT-F and AFfumR-RT-R for *A. fumigatus fumR*, AFhelA-RT-F and AFhelA-RT-R for *A. fumigatus helA*, AFpyr2-RT-F and AFpyr2-RT-R for *A. fumigatus pyr2*, AFact1-RT-F and AFact1-RT-R for *A. fumigatus act1*, AOnrdA-RT-F and AOnrdA-RT-R for *A. oryzae nrdA*, AOkojR-RT-F and AOkojR-RT-R for *A. oryzae kojR*, AOkojA-RT-F and AOkojA-RT-R for *A. oryzae kojA*, AOkojT-RT-F and AOkojT-RT-R for *A. oryzae kojT*, AOipnA-RT-F and AOipnA-RT-R for *A. oryzae ipnA*, and AOactA-RT-F and AOactA-RT-R for *A. oryzae actA* (see Table S2 in the supplemental material). The gene expression levels of *acnA*, *act1*, and *actA* were used to calibrate those of *A. nidulans*, *A. fumigatus*, and *A. oryzae*, respectively.

### Data availability

The data obtained from the RNA-seq and RIP-seq analyses in this study were deposited in the Gene Expression Omnibus under accession number GSE164392.

## ACKNOWLEDGMENTS

We thank the Division of Instrumental Analysis Research Support Center of Kagoshima University for technical support. This study was supported in part by KAKENHI (grant numbers: 16K07672, 18K05394, and 19K05773) and a research grant from Kagoshima University based on a scholarship donation from SUNUS Co., Ltd. C. K. received a Grant-in-Aid for JSPS Research Fellows (grant number: 17J02753).

## REFERENCES

1. Kuehner JN, Pearson EL, Moore C. 2011. Unravelling the means to an end: RNA polymerase II transcription termination. Nat Rev Mol Cell Biol. 12:283–294.

2. Arndt KM, Reines D. 2015. Termination of transcription of short noncoding RNAs by RNA polymerase II. Annu Rev Biochem 84:381–404.

3. Darby MM, Serebreni L, Pan X, Boeke JD, Corden JL. 2012. The *Saccharomyces cerevisiae* Nrd1-Nab3 transcription termination pathway acts in opposition to Ras signaling and mediates response to nutrient depletion. Mol Cell Biol 32:1762–1775.

4. Merran J, Corden JL. 2017. Yeast RNA-binding protein Nab3 regulates genes involved in nitrogen metabolism. Mol Cell Biol 37:e00154–17.

5. Webb S, Hector RD, Kudla G, Granneman S. 2014. PAR-CLIP data indicate that Nrd1-Nab3-dependent transcription termination regulates expression of hundreds of protein coding genes in yeast. Genome Biol 15:R8.

6. Lemay JF, Marguerat S, Larochelle M, Liu X, van Nues R, Hunyadkürti J, Hoque M, Tian B, Granneman S, Bähler J, Bachand F. 2016. The Nrd1-like protein Seb1 coordinates cotranscriptional 3’ end processing and polyadenylation site selection. Genes Dev 30:1558–1572.

7. Wittmann S, Renner M, Watts BR, Adams O, Huseyin M, Baejen C, El Omari K, Kilchert C, Heo DH, Kecman T, Cramer P, Grimes JM, Vasiljeva L. 2017. The conserved protein Seb1 drives transcription termination by binding RNA polymerase II and nascent RNA. Nat Commun 8:14861.

8. Suganuma T, Fujita K, Kitahara K. 2007. Some distinguishable properties between acid-stable and neutral types of alpha-amylases from acid-producing koji. J Biosci Bioeng 104:353–362.

9. Yamada O, Takara R, Hamada R, Hayashi R, Tsukahara M, Mikami S. 2011. Molecular biological researches of Kuro-Koji molds, their classification and safety. J Biosci Bioeng 112:233–237.

10. Futagami T, Mori K, Wada S, Ida H, Kajiwara Y, Takashita H, Tashiro K, Yamada O, Omori T, Kuhara S, Goto M. 2015. Transcriptomic analysis of temperature responses of *Aspergillus kawachii* during barley koji production. Appl Environ Microbiol 81:1353–1363.

11. Kadooka C, Izumitsu K, Onoue M, Okutsu K, Yoshizaki Y, Takamine K, Goto M, Tamaki H, Futagami T. 2019. Mitochondrial citrate transporters CtpA and YhmA are required for extracellular citric acid accumulation and contribute to cytosolic acetyl coenzyme A generation in *Aspergillus luchuensis* mut. *kawachii*. Appl Environ Microbiol 85:e03136–18

12. Nützmann HW, Scazzocchio C, Osbourn A. 2018. Metabolic gene clusters in eukaryotes. Annu Rev Genet 52:159–183.

13. Rokas A, Wisecaver JH, Lind AL. 2018. The birth, evolution and death of metabolic gene clusters in fungi. Nat Rev Microbiol 16:731–744.

14. Steinmetz EJ, Brow DA. 1998. Control of pre-mRNA accumulation by the essential yeast protein Nrd1 requires high-affinity transcript binding and a domain implicated in RNA polymerase II association. Proc Natl Acad Sci U S A 95:6699–6704.

15. Kubicek K, Cerna H, Holub P, Pasulka J, Hrossova D, Loehr F, Hofr C, Vanacova S, Stefl R. 2012. Serine phosphorylation and proline isomerization in RNAP II CTD control recruitment of Nrd1. Genes Dev 26:1891–1896.

16. Vasiljeva L, Kim M, Mutschler H, Buratowski S, Meinhart A. 2008. The Nrd1-Nab3-Sen1 termination complex interacts with the Ser5-phosphorylated RNA polymerase II C-terminal domain. Nat Struct Mol Biol 15:795–804.

17. Franco-Echevarría E, González-Polo N, Zorrilla S, Martínez-Lumbreras S, Santiveri CM, Campos-Olivas R, Sánchez M, Calvo O, González B, Pérez-Cañadillas JM. 2017. The structure of transcription termination factor Nrd1 reveals an original mode for GUAA recognition. Nucleic Acids Res 45:10293– 10305.

18. Conrad NK, Wilson SM, Steinmetz EJ, Patturajan M, Brow DA, Swanson MS, Corden JL. 2000. A yeast heterogeneous nuclear ribonucleoprotein complex associated with RNA polymerase II. Genetics 154:557–571.

19. O’Rourke TW, Loya TJ, Head PE, Horton JR, Reines D. 2015. Amyloid-like assembly of the low complexity domain of yeast Nab3. Prion 9:34–47.

20. Steinmetz EJ and Brow DA. 1996. Repression of gene expression by an exogenous sequence element acting in concert with a heterogeneous nuclear ribonucleoprotein-like protein, Nrd1, and the putative helicase Sen1. Mol Cell Biol 16:6993–7003.

21. Mitsuzawa H, Kanda E, Ishihama A. 2003. Rpb7 subunit of RNA polymerase II interacts with an RNA-binding protein involved in processing of transcripts. Nucleic Acids Res 31:4696–4701.

22. Hashimoto R, Asano K, Tokashiki T, Onji Y, Hirose-Yasumoto M, Takara R, Toyosato T, Yoshino A, Ikehata M, Ying L, Kumeda Y, Yokoyama K, Takahashi H. 2013. Mycotoxin production and genetic analysis of *Aspergillus niger* and related species including the Kuro-Koji mold. Mycotoxins 63:179–186.

23. Yamada O, Machida M, Hosoyama A, Goto M, Takahashi T, Futagami T, Yamagata Y, Takeuchi M, Kobayashi T, Koike H, Abe K, Asai K, Arita M, Fujita N, Fukuda K, Higa KI, Horikawa H, Ishikawa T, Jinno K, Kato Y, Kirimura K, Mizutani O, Nakasone K, Sano M, Shiraishi Y, Tsukahara M, Gomi K. 2016. Genome sequence of *Aspergillus luchuensis* NBRC 4314. DNA Res 23:507–515.

24. Ramón D, Carramolino L, Patiño C, Sánchez F, Peñalva MA. 1987. Cloning and characterization of the isopenicillin N synthetase gene mediating the formation of the β-lactam ring in *Aspergillus nidulans*. Gene 57:171–181.

25. Yu JH, Butchko RA, Fernandes M, Keller NP, Leonard TJ, Adams TH. 1996. Conservation of structure and function of the aflatoxin regulatory gene *aflR* from *Aspergillus nidulans* and *A. flavus*. Curr Genet 29:549–555.

26. Keller NP, Kantz NJ, Adams TH. 1994. *Aspergillus nidulans verA* is required for production of the mycotoxin sterigmatocystin. Appl Environ Microbiol 60:1444– 1450.

27. Brown DW, Yu JH, Kelkar HS, Fernandes M, Nesbitt TC, Keller NP, Adams TH, Leonard TJ. 1996. Twenty-five coregulated transcripts define a sterigmatocystin gene cluster in *Aspergillus nidulans*. Proc Natl Acad Sci U S A 93:1418–1422.

28. Dhingra S, Lind AL, Lin HC, Tang Y, Rokas A, Calvo AM. 2013. The fumagillin gene cluster, an example of hundreds of genes under *veA* control in *Aspergillus fumigatus*. PLoS One 8:e77147.

29. Lv JM, Hu D, Gao H, Kushiro T, Awakawa T, Chen GD, Wang CX, Abe I, Yao XS. 2017. Biosynthesis of helvolic acid and identification of an unusual C-4-demethylation process distinct from sterol biosynthesis. Nat Commun 8:1644.

30. Itoh T, Tokunaga K, Matsuda Y, Fujii I, Abe I, Ebizuka Y, Kushiro T. 2010. Reconstitution of a fungal meroterpenoid biosynthesis reveals the involvement of a novel family of terpene cyclases. Nat Chem 2:858–864.

31. Terabayashi Y, Sano M, Yamane N, Marui J, Tamano K, Sagara J, Dohmoto M, Oda K, Ohshima E, Tachibana K, Higa Y, Ohashi S, Koike H, Machida M. 2010. Identification and characterization of genes responsible for biosynthesis of kojic acid, an industrially important compound from *Aspergillus oryzae*. Fungal Genet Biol 47:953–961.

32. Keller NP. 2018. Fungal secondary metabolism: regulation, function and drug discovery. Nat Rev Microbiol 17:167–180.

33. Lind AL, Lim FY, Soukup AA, Keller NP, Rokas A. 2018. An LaeA- and BrlA-dependent cellular network governs tissue-specific secondary metabolism in the human pathogen Aspergillus fumigatus. mSphere 3:e00050–18.

34. Bok JW, Keller NP. 2004. LaeA, a regulator of secondary metabolism in *Aspergillus* spp. Eukaryot Cell 3:527–535.

35. Oda K, Kobayashi A, Ohashi S, Sano M. 2011. *Aspergillus oryzae laeA* regulates kojic acid synthesis genes. Biosci Biotechnol Biochem 75:1832–1834.

36. Kawauchi M, Nishiura M, Iwashita K. 2013. Fungus-specific sirtuin HstD coordinates secondary metabolism and development through control of LaeA. Eukaryot Cell 12:1087–1096.

37. Vasiljeva L, Kim M, Terzi N, Soares LM, Buratowski S. 2008. Transcription termination and RNA degradation contribute to silencing of RNA polymerase II transcription within heterochromatin. Mol Cell 29:313–323.

38. Parsa JY, Boudoukha S, Burke J, Homer C, Madhani HD. 2018. Polymerase pausing induced by sequence-specific RNA-binding protein drives heterochromatin assembly. Genes Dev 32:953–964.

39. Niu J, Arentshorst M, Nair PD, Dai Z, Baker SE, Frisvad JC, Nielsen KF, Punt PJ, Ram AF. 2015. Identification of a classical mutant in the industrial host *Aspergillus niger* by systems genetics: LaeA is required for citric acid production and regulates the formation of some secondary metabolites. G3 (Bethesda) 6:193–204.

40. Linde T, Zoglowek M, Lübeck M, Frisvad JC, Lübeck PS. 2016. The global regulator LaeA controls production of citric acid and endoglucanases in *Aspergillus carbonarius*. J Ind Microbiol Biotechnol 43:1139–1147.

41. Kadooka C, Nakamura E, Mori K, Okutsu K, Yoshizaki Y, Takamine K, Goto M, Tamaki H, Futagami T. 2020. LaeA controls citric acid production through regulation of the citrate exporter-encoding *cexA* gene in *Aspergillus luchuensis* mut. *kawachii*. Appl Environ Microbiol 86:e01950–19.

42. Kadooka C, Onitsuka S, Uzawa M, Tashiro S, Kajiwara Y, Takashita H, Okutsu K, Yoshizaki Y, Takamine K, Goto M, Tamaki H, Futagami T. 2016. Marker recycling system using the *sC* gene in the white koji mold, Aspergillus luchuensis mut. kawachii. J Gen Appl Microbiol 62:160–163.

43. Mizutani O, Kudo Y, Saito A, Matsuura T, Inoue H, Abe K, Gomi K. 2008. A defect of LigD (human Lig4 homolog) for nonhomologous end joining significantly improves efficiency of gene-targeting in *Aspergillus oryzae*. Fungal Genet Biol 45:878–889.

44. Nakamura E, Kadooka C, Okutsu K, Yoshizaki Y, Takamine K, Goto M, Tamaki H, Futagami T. 2021. Citrate exporter enhances both extracellular and intracellular citric acid accumulation in the koji fungi *Aspergillus luchuensis* mut. *kawachii* and *Aspergillus oryzae*. J Biosci Bioeng 131:68–76.

45. Barratt RW, Johnson GB, Ogata WN. 1965. Wild-type and mutant stocks of *Aspergillus nidulans*. Genetics 52:233–246.

46. Meyer V, Wanka F, van Gent J, Arentshorst M, van den Hondel CA, Ram AF. 2011. Fungal gene expression on demand: an inducible, tunable, and metabolism-independent expression system for *Aspergillus niger*. Appl Environ Microbiol 77:2975–2983.

47. Kubodera T, Yamashita N, Nishimura A. 2002. Transformation of *Aspergillus* sp. and *Trichoderma reesei* using the pyrithiamine resistance gene (*ptrA*) of *Aspergillus oryzae*. Biosci Biotechnol Biochem 66:404–406.

48. Liu HL, Osmani AH, Ukil L, Son S, Markossian S, Shen KF, Govindaraghavan M, Varadaraj A, Hashmi SB, De Souza CP, Osmani SA. 2010. Single-step affinity purification for fungal proteomics. Eukaryot Cell 9:831–833.

49. Yang L, Ukil L, Osmani A, Nahm F, Davies J, De Souza CP, Dou X, Perez-Balaguer A, Osmani SA. 2004. Rapid production of gene replacement constructs and generation of a green fluorescent protein-tagged centromeric marker in *Aspergillus nidulans*. Eukaryot Cell 3:1359–1362

50. Miyamoto A, Kadooka C, Mori K, Tagawa Y, Okutsu K, Yoshizaki Y, Takamine K, Goto M, Tamaki H, Futagami T. 2020. Sirtuin SirD is involved in alpha-amylase activity and citric acid production in *Aspergillus luchuensis* mut. *kawachii* during a solid-state fermentation process. J Biosci Bioeng 129:454–466.

51. Futagami T, Mori K, Yamashita A, Wada S, Kajiwara Y, Takashita H, Omori T, Takegawa K, Tashiro K, Kuhara S, Goto M. 2011. Genome sequence of the white koji mold *Aspergillus kawachii* IFO 4308, used for brewing the Japanese distilled spirit shochu. Eukaryot Cell 10:1586–1587.

52. Bolger AM, Lohse M, Usadel B. 2014. Trimmomatic: a flexible trimmer for Illumina sequence data. Bioinformatics 30:2114–2120.

53. Martin M. 2011. Cutadapt removes adapter sequences from high-throughput sequencing reads. EMBnet J 17:10

54. Dobin A, Gingeras TR. 2016. Optimizing RNA-Seq Mapping with STAR. Methods Mol Biol 1415:245–262.

55. Liao Y, Smyth GK, Shi W. 2014. featureCounts: an efficient general purpose program for assigning sequence reads to genomic features. Bioinformatics 30:923– 930.

56. Love MI, Huber W, Anders S. 2014. Moderated estimation of fold change and dispersion for RNA-seq data with DESeq2. Genome Biol 15:550.

57. Si Y, Liu P, Li P, Brutnell TP. 2014. Model-based clustering for RNA-seq data. Bioinformatics 30:197–205.

58. Cerqueira GC, Arnaud MB, Inglis DO, Skrzypek MS, Binkley G, Simison M, Miyasato SR, Binkley J, Orvis J, Shah P, Wymore F, Sherlock G, Wortman JR (2014). The *Aspergillus* Genome Database: multispecies curation and incorporation of RNA-Seq data to improve structural gene annotations. Nucleic Acids Res 42:D705–10.

59. Hong SB, Lee M, Kim DH, Varga J, Frisvad JC, Perrone G, Gomi K, Yamada O, Machida M, Houbraken J, Samson RA. 2013. *Aspergillus luchuensis*, an industrially important black *Aspergillus* in East Asia. PLoS One 8:e63769.

60. Hong SB, Yamada O, Samson RA. 2014. Taxonomic re-evaluation of black koji molds. Appl Microbiol Biotechnol 98:555–561.

